# *Pseudomonas aeruginosa* Can Inhibit Growth of Streptococcal Species via Siderophore Production

**DOI:** 10.1101/307488

**Authors:** Jessie E. Scott, Kewei Li, Laura M. Filkins, Bin Zhu, Sherry L. Kuchma, Joseph D. Schwartzman, George A. O’Toole

## Abstract

Cystic Fibrosis (CF) is a genetic disease that causes patients to accumulate thick, dehydrated mucus in the lung and develop chronic, polymicrobial infections due to reduced mucociliary clearance. These chronic polymicrobial infections and subsequent decline in lung function are significant factors in the morbidity and mortality of CF. *Pseudomonas aeruginosa* and *Streptococcus* spp. are among the most prevalent organisms in the CF lung; the presence of *P. aeruginosa* correlates with lung function decline and the *Streptococcus milleri* group (SMG), a subgroup of the viridans streptococci, is associated with exacerbations in patients with CF. Here we characterize the interspecies interactions that occur between these two genera. We demonstrated that multiple *P. aeruginosa* laboratory strains and clinical CF isolates promote the growth of multiple SMG strains and oral streptococci in an *in vitro* coculture system. We investigated the mechanism by which *P. aeruginosa* enhances growth of streptococci by screening for mutants of *P. aeruginosa* PA14 unable to enhance *Streptococcus* growth, and we identified the *P. aeruginosa pqsL::TnM* mutant, which failed to promote growth of *S. constellatus* and *S. sanguinis*. Characterization of the *P. aeruginosa* Δ*pqsL* mutant revealed that this strain cannot promote *Streptococcus* growth. Our genetic data and growth studies support a model whereby the *P. aeruginosa* Δ*pqsL* mutant overproduces siderophores, and thus likely outcompetes *Streptococcus sanguinis* for limited iron. We propose a model whereby competition for iron represents one important means of interaction between *P. aeruginosa* and *Streptococcus* spp.

**Importance:** Cystic fibrosis (CF) lung infections are increasingly recognized for their polymicrobial nature. These polymicrobial infections may alter the biology of the organisms involved in CF-related infections, leading to changes in growth, virulence and/or antibiotic tolerance, and could thereby affect patient health and response to treatment. In this study, we demonstrate interactions between *P. aeruginosa* and streptococci using a coculture model, and show that one interaction between these microbes is likely competition for iron. Thus, these data indicate that one CF pathogen may influence the growth of another and add to our limited knowledge of polymicrobial interactions in the CF airway.

## Introduction

Cystic fibrosis (CF) is a genetic disease caused by a defect in the cystic fibrosis transmembrane conductance regulator (1), which leads to reduced mucociliary clearance in the lungs of these patients (2). Due to this reduced mucociliary clearance, bacteria colonize the lungs of patients with CF and establish chronic, polymicrobial infections that cause increased inflammation and respiratory function decline (2). Recent studies have demonstrated that the microbiota in the lungs form polymicrobial biofilms, and that mixed bacterial biofilm populations can affect antibiotic tolerance and bacterial virulence (3, 4).

The *Streptococcus milleri* group (SMG), which is composed of three species (*S. anginosus, S. constellatus*, and *S. intermedius*), has been isolated from sputum samples of patients with CF. When these microbes are the numerically dominant species in the lung, these organisms correlate with exacerbation in patients with CF (5-8). In contrast, previous research from our laboratory (9) and two other groups (10, 11) demonstrated that increased relative abundance of *Streptococcus* spp. within the CF lung microbiome correlates with better lung function and clinical stability. Together these data indicate a possible complex relationship between *Streptococcus* spp. and the host, and *Streptococcus* spp. and the other microbes in the CF airway.

*Pseudomonas aeruginosa* is the dominant microorganism (>50% relative abundance) in the lungs of ~45% of adults patients with CF (12), is cultured from >80% of these patients (13), and is the predominant microbe in the lung at end stage disease (14). *P. aeruginosa* and streptococci have been found to co-colonize CF patients (5, 6, 8, 15, 16), but the polymicrobial interactions that occur between these organisms are not well studied. Previous studies investigating interactions between *P. aeruginosa* and *Streptococcus* spp. demonstrated that *Streptococcus* spp. can influence production of *P. aeruginosa* virulence factors such as rhamnolipids, elastase, and phenazines (7, 17–20), and can suppress *P. aeruginosa* growth through hydrogen peroxide production (17) and production of reactive nitrogenous intermediates (21, 22). Conversely, *P. aeruginosa* was found to influence the growth (17–20, 23, 24) and biofilm formation (24, 25) of *Streptococcus* spp. Work from our lab demonstrated that *P. aeruginosa* PA14 produces the surfactants β-hydroxyalkanoyl-β-hydroxyalkanoic acids (HAAs) and monorhamnolipids which caused a 6-fold reduction in *S. constellatus* 7155 biofilm formation in coculture (24). The surfactant-induced biofilm suppression was relieved when *P. aeruginosa* and *S. constellatus* 7155 were cocultured in the presence of tobramycin, an antibiotic used for maintenance therapy by patients with CF. We determined that tobramycin suppressed *P. aeruginosa* production of HAAs and monorhamnolipids, and that in the presence of tobramycin, *P. aeruginosa* can enhance *S. constellatus* 7155 growth on a CF-derived bronchial epithelial cell (CFBE) monolayer (24). These data indicate that *P. aeruginosa* can both positively and negatively impact cocultured microbes, including *Streptococcus* spp., and the interaction between the microbes can be influenced by environmental and/or clinical context.

In this study, we investigate the ability of *P. aeruginosa* to influence *Streptococcus* growth in our *in vitro* coculture system. We demonstrate that multiple *P. aeruginosa* strains and clinical isolates can enhance the growth of multiple *Streptococcus* spp. We used a candidate gene approach and a genetic screen to identify *P. aeruginosa* mutants that were unable to support *Streptococcus* growth, and found a single mutant of *P. aeruginosa* that no longer enhances growth of streptococci. We found that the *P. aeruginosa* Δ*pqsL* mutant suppressed *S. sanguinis* growth, likely via a mechanism that involves siderophore overproduction and thus iron sequestration. These data indicate that competition for iron can impact this polymicrobial interaction.

## Results

### *P. aeruginosa* promotes streptococcal growth in a coculture system

We reported previously that *P. aeruginosa* can enhance viable *S. constellatus* 7155 cell number when grown as a coculture on CF-derived bronchial epithelial (CFBE) cells (24). We first sought to recapitulate the finding that *P. aeruginosa* promotes the *Streptococcus* biofilm population by using the model organism *S. sanguinis* SK36 in coculture conditions in the absence of CFBE cells. We used *P. aeruginosa* PAO1 and *S. sanguinis* SK36 for these experiments because both are sequenced strains (26–28) with available genetic mutant libraries (29–31). This simplified coculture system allowed us to test the interaction between *P. aeruginosa* and streptococci without confounding factors contributed by the CFBE cells.

To test the hypothesis that *S. sanguinis* SK36 viable cell number increases in coculture with *P. aeruginosa* PAO1 in absence of CFBE cells, we grew *P. aeruginosa* PAO1 and *S. sanguinis* SK36 in coculture in the wells of a plastic culture dish in the minimal medium MEM tissue culture medium containing glucose. We observed that the number of viable *S. sanguinis* SK36 in a biofilm was enhanced 100-1055-fold by coculture with *P. aeruginosa* PAO1 compared to *S. sanguinis* SK36 grown as a monoculture in MEM (Fig. 1A, see also Fig. S1A). *P. aeruginosa* PAO1 biofilm growth was not significantly affected by coculture with *S. sanguinis* SK36 (Fig. S1B). These data also indicate that the enhancement of the *S. sanguinis* SK36 population in a biofilm by *P. aeruginosa* PAO1 does not require the CFBE cells.

**Figure 1.**
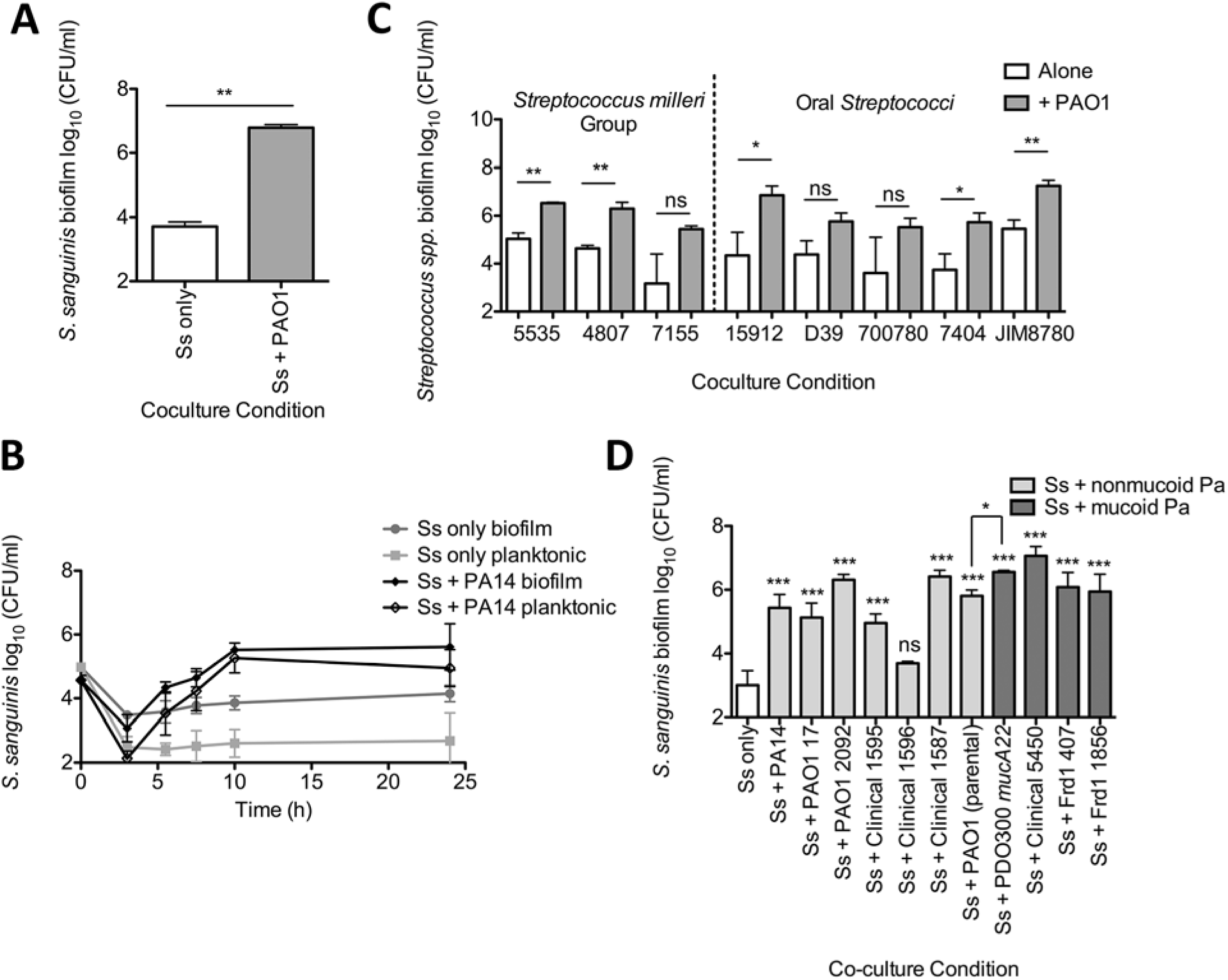
*P. aeruginosa* enhanced growth of *Streptococcus* spp. in coculture. (A to D) Coculture assays were conducted to investigate streptococcal growth when cocultured with *P. aeruginosa*. (A) *S. sanguinis* SK36 (Ss) cocultured with *P. aeruginosa* PAO1. (B) Coculture growth kinetics of *S. sanguinis* with *P. aeruginosa* PA14 were investigated. The time points each represent the average of three biological replicates with three technical replicates. The error bars indicate SD. (C) A representative group of clinical and reference *Streptococcus* strains were tested in coculture with *P. aeruginosa* PAO1 (see Fig. S3A for all *Streptococcus* spp. strains tested). In this panel, each strain is denoted by their strain number: *S. anginosus* 5535, *S. intermedius* 4807, *S. constellatus* 7155, *S. parasanguinis* ATCC15912, *S. pneumoniae* D39, *S. peroris* ATCC700780, *S. oralis* 7404, and *S. salivarius* JIM8780. (D) *S. sanguinis* SK36 was tested in coculture with multiple *P. aeruginosa* clinical and laboratory strains. (A, C, D) Each column represents the average of three biological replicates with three technical replicates. The error bars indicate the standard deviation. ns, not significant, ^*^, P < 0.05, **, P < 0.01, and ^***^, P < 0.001 by paired two-tailed student’s t-test (A, C), repeated measures one-way analysis of variance (ANOVA) with Tukey’s multiple comparisons posttest (D), and paired two-tailed student’s t-test between Ss + PAO1 DHL08 and Ss + PAO1 mucA22 (D). The corresponding graphs depicting *P. aeruginosa* growth in these assays can be found in Figures S1-4.

### *P. aeruginosa* enhances growth of *S. sanguinis* SK36

We considered two models of polymicrobial interaction that may be enhancing viable *S. sanguinis* SK36 cells in the biofilm when grown in coculture with *P. aeruginosa. P. aeruginosa* might promote *Streptococcus* adhesion and biofilm formation, or *P. aeruginosa* may promote streptococcal growth. To distinguish between these models, we conducted a time course experiment with *P. aeruginosa* PA14, *P. aeruginosa* PAO1, and *S. sanguinis* SK36. If *P. aeruginosa* was promoting adhesion of *S. sanguinis* SK36 cells rather than growth, we predict that we would detect more *S. sanguinis* SK36 in the biofilm and fewer planktonic cells, but total cell number would not increase compared to *S. sanguinis* SK36 monoculture. In contrast, if *P. aeruginosa* were enhancing *S. sanguinis* SK36 growth, then both total biofilm and planktonic *S. sanguinis* SK36 populations should increase in coculture compared to *S. sanguinis* SK36 monoculture. As demonstrated by the increased *S. sanguinis* SK36 biofilm and planktonic cells recovered from coculture compared to monoculture, *P. aeruginosa* appears to promote the growth of *S. sanguinis* SK36 (Fig. 1B, and Fig. S2A and S2B), thus accounting for the increased population of *S. sanguinis* SK36 biofilm cells.

### Multiple *P. aeruginosa* strains enhance the growth of multiple streptococci

Based on current evidence that multiple *Streptococcus* species inhabit the CF lung (8, 32) and influence patient health (5, 6, 8–11, 15, 16), we sought to determine whether the observed enhancement of *Streptococcus* viable counts in coculture with *P. aeruginosa* may be more broadly generalized to other streptococci, including the *Streptococcus milleri* group (SMG), which has been implicated in CF-related exacerbations (5, 6, 8, 15). To assess the ability of *P. aeruginosa* to promote multiple *Streptococcus* spp., we cocultured *P. aeruginosa* PAO1 with 6 SMG isolates and 8 oral *Streptococcus* spp. Figure 1C depicts a representative strain of each streptococcal species assayed, and shows the biofilm population obtained from monoculture and coculture with *P. aeruginosa* PAO1, respectively (see also Fig. S3A for all 14 strains tested). *P. aeruginosa* PAO1 growth was not significantly affected by coculture with any of the *Streptococcus* spp. tested (Fig. S3B).

We found that *P. aeruginosa* PAO1 significantly enhanced the growth of one of the two *S. anginosus*, two of two *S. intermedius*, and neither of the two *S. constellatus* strains tested. Additionally, of the oral *Streptococcus* spp. tested, *P. aeruginosa* PAO1 significantly promoted the growth of one of the two *S. oralis*, the one *S. parasanguinis*, and one of the three *S. salivarius* isolates, but not the *S. pneumoniae* or *S. peroris* isolates tested (Fig. 1C and Fig. S3A). While not every *Streptococcus* isolate tested demonstrated significant increase in viable population recovered from the coculture, most species tested exhibited a trend toward increased growth when cocultured with *P. aeruginosa* PAO1. These data suggest that *P. aeruginosa* may be promoting *Streptococcus* growth through a pathway that affects many *Streptococcus* species.

Next we assessed whether multiple *P. aeruginosa* clinical and laboratory strains could promote the growth of *S. sanguinis*. Additionally, given that *S. parasanguinis* was found to bind extracellular alginate produced by mucoid *P. aeruginosa* strains (25), we tested whether mucoid or nonmucoid *P. aeruginosa* could better promote growth in our coculture system. We cocultured *S. sanguinis* SK36 with seven nonmucoid *P. aeruginosa* and four mucoid *P. aeruginosa* laboratory and clinical strains, and observed a significant growth enhancement of *S. sanguinis* SK36 by ten out of eleven *P. aeruginosa* strains tested in coculture biofilms (Fig. 1D) and planktonic growth (Fig. S4A). The growth of all tested *P. aeruginosa* strains was not affected by coculture with *S. sanguinis* SK36 (Fig. S4B, S4C).

Additionally, *P. aeruginosa* PAO1 (parental) and *P. aeruginosa* PDO300 *mucA22* are isogenic nonmucoid and mucoid strains, respectively. We found a significant enhancement in viable *S. sanguinis* SK36 biofilm cells recovered from coculture with *P. aeruginosa* PDO300 *mucA22* compared to *P. aeruginosa* PAO1 (Fig. 1D), suggesting that mucoid *P. aeruginosa* strains may better enhance *Streptococcus* growth. Additionally, these mucoid *P. aeruginosa* strains showed among the most robust promotion of viable counts when cocultured with *Streptococcus*.

To extend our findings here, we next tested the growth enhancing capability of *P. aeruginosa* PA14 in rich medium, using both lysogeny broth (LB) and Todd Hewitt broth supplemented with 0.5% yeast extract (THY) and found that *P. aeruginosa* PA14 is able to significantly enhance *S. sanguinis* SK36 growth in these conditions (Fig. S4D). These data suggest that *P. aeruginosa* may be providing nutrients to *S. sanguinis* SK36 in minimal medium, given that *S. sanguinis* SK36 growth in rich medium monoculture reaches a level comparable to that of coculture in minimal medium. However, there is still a significant increase in *S. sanguinis* SK36 growth when in coculture with *P. aeruginosa* PA14 in rich medium, perhaps indicating non-nutritional mechanisms of growth enhancement by *P. aeruginosa*.

In summary, we have demonstrated that our minimal medium coculture assay using a plastic substratum can recapitulate our prior observation that *P. aeruginosa* promotes streptococcal growth on airway cells. This observation extends to coculture assays in rich medium. Furthermore, we were able to determine that *P. aeruginosa* is likely promoting *Streptococcus* growth rather than increasing the biofilm population via enhanced adherence. The *Streptococcus* growth-enhancement phenotype occurred among most oral streptococci tested, and the majority of *P. aeruginosa* clinical and laboratory strains are capable of promoting *Streptococcus* growth, which lends support to the idea that these interactions are common among these two genera.

## Known *P. aeruginosa* virulence pathways are not involved in the *Streptococcus* growth-promoting phenotype

*P. aeruginosa* has many well characterized virulence factors that have been demonstrated to impact polymicrobial interactions, including pathways for quorum sensing (33), biofilm formation, and the production of secreted molecules such as phenazines (34, 35), siderophores (3, 36), alginate (37), and rhamnolipids (24). We hypothesized that one or more of these virulence factors might be altering *Streptococcus* growth in our system. To test this idea, we utilized a candidate genetic approach to assess whether any of these virulence pathways may be involved in the observed growth-enhancing phenotype. We cocultured *P. aeruginosa* PA14 mutants in each of the above pathways with *S. constellatus* 7155 as a model streptococcal strain know to positively respond to *P. aeruginosa* growth enhancement (24, 38), and assessed whether any of these mutants lost the ability to enhance *S. constellatus* 7155 growth. We found that none of the pathways tested were involved in enhancement of *S. constellatus* 7155 growth (Table S1).

Given that KatA has been found in the supernatant of *P. aeruginosa* cultures (38–40) we also constructed *P. aeruginosa* PA14 Δ*katA*, Δ*katB*, and Δ*katΔkatB* mutant strains in order to test the hypothesis that extracellular *P. aeruginosa* catalase is enhancing *S. sanguinis* SK36 growth by degrading hydrogen peroxide produced by *S. sanguinis* SK36. It has previously been reported that *S. sanguinis* and other oral streptococci can inhibit *P. aeruginosa* growth through hydrogen peroxide production (17, 21, 22), and that the hydrogen peroxide produced by oral streptococci plays an important role in growth inhibition, eDNA release, and biofilm formation within the oral microbiome (41, 42). We chose to mutate the *katA and katB* genes and not the *katE* gene because previous reports indicate that KatA is the major catalase utilized by *P. aeruginosa*, and that KatB can partially recover hydrogen peroxide resistance in the absence of KatA (38, 43). KatE was not demonstrated to play a role in alleviating hydrogen peroxide stress (43). *S. sanguinis* SK36 did not demonstrate reduced growth in coculture with the *P. aeruginosa* PA14 Δ*katA*, Δ*katB*, or Δ*katA*Δ*katB* mutant strains compared to wild-type *P. aeruginosa* PA14, indicating that catalase is not playing a role in the *Streptococcus* growth enhancement phenotype (Table S1 and Fig. S5A). The *P. aeruginosa* Δ*katA* mutant displays a slight, but significant growth defect in the coculture compared to wild-type *P. aeruginosa* PA14 in coculture, and the Δ*katB* mutant displays a modest, but significant growth defect in monoculture compared to *P. aeruginosa* PA14 in coculture (Fig. S5B), but these strains still stimulate *S. sanguinis* SK36 growth to the level observed for wild-type *P. aeruginosa*. Taken together, these data suggest known virulence factors, on their own, do not contribute to *P. aeruginosa-mediated* growth enhancement of *Streptococcus* spp.

### Screening the *P. aeruginosa* PA14NR Set for *P. aeruginosa* PA14 transposon insertion mutant strains that do not support *S. constellatus* growth

The Ausubel lab reported a nonredundant library of *P. aeruginosa* PA14 transposon insertion mutants (PA14NR Set) containing 5,459 transposon insertion mutant strains with mutations in 4,596 genes (44). Each of these *P. aeruginosa* PA14 transposon mutant strains were tested in coculture with *S. constellatus* 7155 (Fig. 2A). Of the 5,459 mutant strains in the library, 48 strains were unable to promote *S. constellatus* 7155 growth in two replicate experiments (Table S2). Two of these 48 mutants were eliminated when we tested available deletion mutants as the deletion mutant strains did not recapitulate the phenotype of the transposon mutation (not shown). The remaining 46 transposon mutants (Table S2) were tested in our standard coculture assay with *S. sanguinis* SK36 to determine which *P. aeruginosa* PA14 transposon mutants are unable to enhance *Streptococcus* growth in a second strain. 44 of the 46 *P. aeruginosa* PA14 transposon mutants were capable of enhancing growth of *S. sanguinis* SK36, and thus were unlikely involved in a general pathway for enhancing growth of *Streptococcus*. We found that two transposon mutants were unable to promote either *S. constellatus* 7155 or *S. sanguinis* SK36 growth: *P. aeruginosa pqsL::TnM*and *P. aeruginosa dbpA::TnM*.

**Figure 2.**
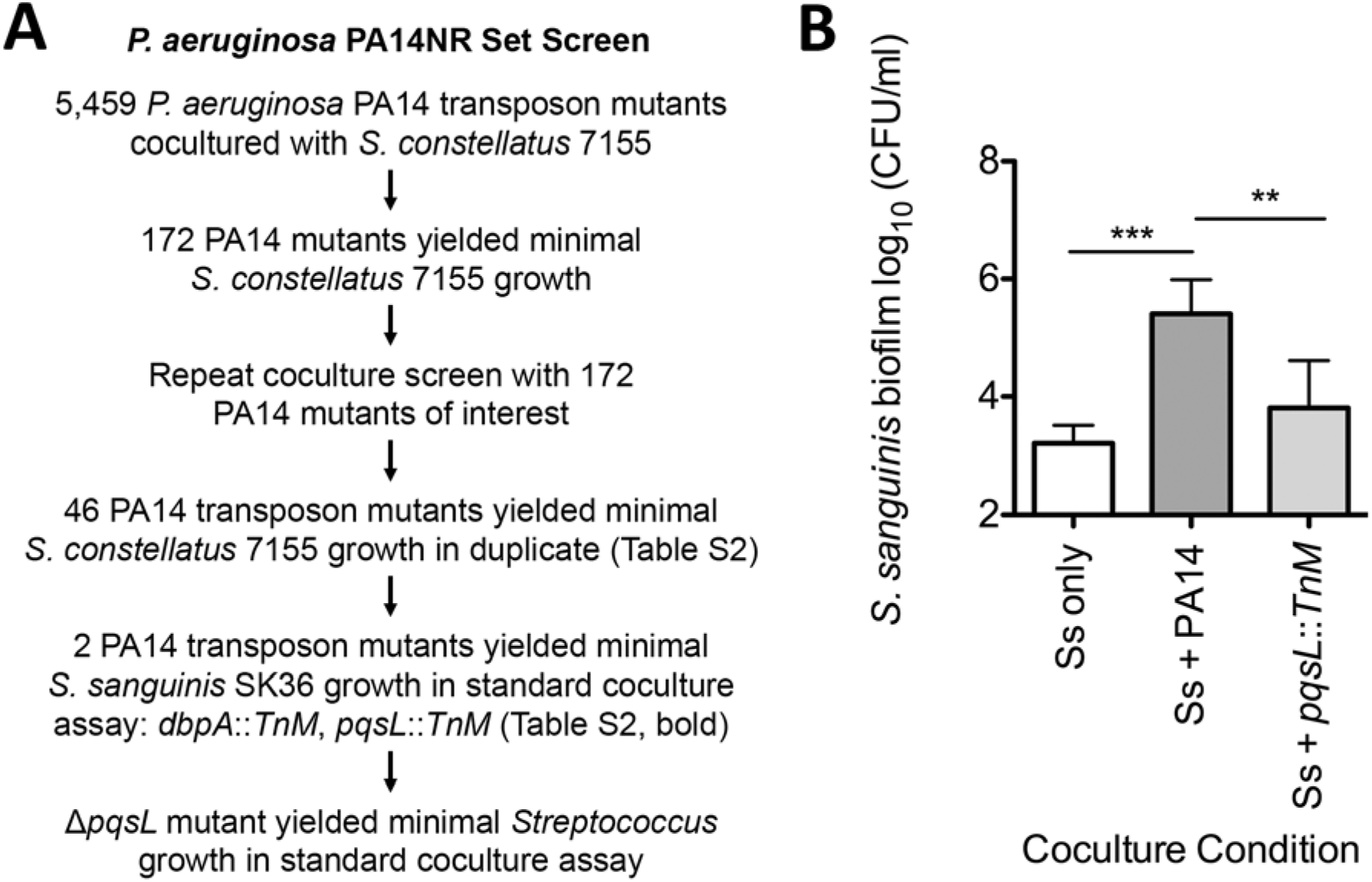
Screening for *P. aeruginosa* PA14 mutants altered in interaction with *Streptococcus*. (A) An overview of the *P. aeruginosa* PA14NR Set transposon mutant screen used to identify *P. aeruginosa* transposon insertion mutants that can no longer enhance *S. constellatus* 7155 growth, and the number of *P. aeruginosa* PA14 transposon insertion mutants identified in each step. (B) Coculture between a mutant strain identified in the screen, *P. aeruginosa pqsL::TnM*, and *S. sanguinis* SK36. Each bar represents the average of three biological replicates, each with three technical replicates. Error bars represent SD. **, P < 0.01 and ***, P < 0.001 by repeated measures ANOVA with Tukey’s multiple comparison posttest. *P. aeruginosa* growth data from this assay can be found in Figure S7.

The *dbpA* gene codes for the RNA helicase DbpA, which has been demonstrated to play a role in the formation of the 50S ribosomal subunit in *Escherichia* coli (45). *E. coli* is able to compensate for Δ*dbpA* deletions in forming the 50S ribosomal subunit, as described previously (46); an inability to form the 50S ribosomal subunit would otherwise cause a lethal protein synthesis defect, and a dominant negative *dbpA* mutation is necessary to observe a defect in DbpA function in *E. coli*. We built and assayed the *P. aeruginosa* PA14 Δ*dbpA* mutant strain and found no significant defect in *S. sanguinis* SK36 growth enhancement (Fig. S6A) or in *P. aeruginosa* growth (Fig. S6B), and thus did not pursue further study of this mutant.

We previously studied the effects of the *Pseudomonas* quinolone signal pathway (*pqs*) on interactions between *P. aeruginosa* and *Staphylococcus aureus*, including the utilization of 2-heptyl-4-hydroxyquinoline-W-oxide (HQNO), a respiratory chain inhibitor, to drive *S. aureus* to fermentative metabolism (3, 37, 47). As was observed for *S. constellatus* 7155, there was a significant reduction in the ability of *P. aeruginosa pqsL::TnM* mutant to support *S. sanguinis* SK36 growth compared to the wild-type *P. aeruginosa* PA14 (Fig. 2B). There was no detectable growth defect of the *pqsL::TnM* mutant strain compared to wild-type *P. aeruginosa* PA14 in our assay condition (Fig. S7). We chose to focus on the *P. aerugionsa pqsL::TnM* mutant for the remainder of our study.

### The *P. aeruginosa* Δ*pqsL* mutant has a defect in *Streptococcus* growth enhancement.

The PQS pathway involves the production of multiple 4-hydroxy-2-alkylquinolones (HAQs) and begins with anthranilic acid, which is converted to intermediates of unknown structure by the enzymes PqsA and PqsD (Fig. 3A). These unknown intermediates can then be converted into HQNO by PqsL, our gene of interest, or 4-hydroxy-2-heptylquinoline (HHQ) by PqsB and PqsC. HHQ can then be converted into 3,4-dihydroxy-2-heptylquinoline (PQS) by PqsH (48–50). MvfR (also known as PqsR) is the transcriptional regulator that is activated by HHQ and PQS and positively regulates the transcription of operons involved in PQS production, and the LasR, and RhlR quorum sensing pathways, as well as the operons required for production of the siderophores pyoverdine and pyochelin (51, 52). MvfR/PqsR and PQS have also been demonstrated to indirectly increase expression of the phenazine pyocyanin (51).

**Figure 3.**
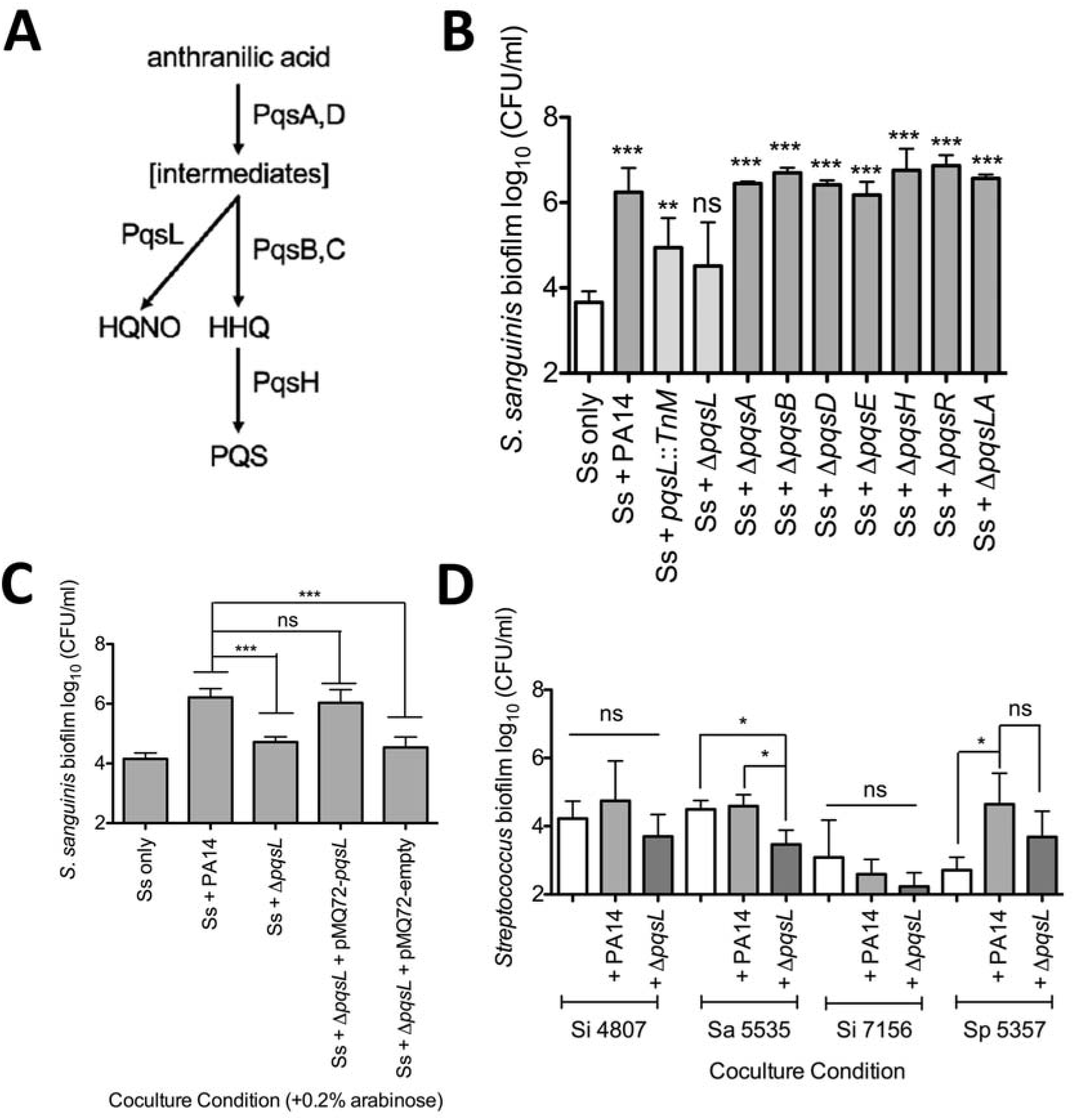
The *P. aeruginosa* Δ*pqsL* mutant inhibits *Streptococcus* growth. (A) The PQS biosynthetic pathway and the enzymes that catalyze each step are shown. (B) Coculture of *S. sanguinis* SK36 with wild-type *P. aeruginosa* PA14 and *P. aeruginosa* PA14 mutant strains lacking each enzyme in the PQS biosynthetic pathway. (C) Coculture of *S. sanguinis* SK36 with *P. aeruginosa* PA14, the Δ*pqsL* mutant, its complement pMQ72-pqsL and the vector control pMQ72 in the presence of 0.2% arabinose. (D) Coculture of representative streptococci from Figure 1 in coculture with the wild-type *P. aeruginosa* PA14 and the Δ*pqsL* mutant strain. In each panel, bars represent the average of three biological replicates, each with at least three technical replicates. Error bars indicate SD. ns, not significant, ^*^, P < 0.05, **, P < 0.01, and ^***^, P < 0.001 by repeated measures ANOVA with Dunnett’s multiple comparisons posttest with *Streptococcus-only* as the control condition (B) and repeated measures ANOVA with Tukey’s multiple comparisons posttest (C and D). The corresponding *P. aeruginosa* growth data for these experiments can be found in Figures S8-10.

We considered two different mechanisms that could explain why the growth of *S. sanguinis* SK36 is no longer promoted by coculture with the *P. aeruginosa* PA14 *pqsL::TnM*. We hypothesized that either the *pqsL::TnM* strain may no longer be able to promote *S. sanguinis* SK36 growth, or the loss of PqsL function resulted in a *P. aeruginosa* strain that reduced *S. sanguinis* SK36 viability. To distinguish between these hypotheses, we assessed the *Streptococcus* growth enhancement capabilities of *P. aeruginosa* PA14 deletion mutants in the *pqs* pathway when grown in coculture with *S. sanguinis* SK36. We found that the Δ*pqsL* mutant was the only mutant in the *pqs* pathway that was unable to promote *S. sanguinis* SK36 growth (Fig. 3B, and Fig. S8A for *S. sanguinis* SK36 planktonic growth, Fig. S8B-C for *P. aeruginosa* biofilm and planktonic growth).

We complemented the Δ*pqsL* strain with an arabinose inducible pMQ72-pqsL construct and demonstrated a significant increase in viable *S. sanguinis* SK36 biofilm cells recovered when the complemented strain was induced with 0.2% arabinose (Fig. 3C, and Fig. S9A for planktonic growth); there was no significant difference between wild-type *P. aeruginosa* PA14 and the complemented ΔpqsL/pMQ72-pqsL strain. Additionally, there was no significant difference in *P. aeruginosa* biofilm and planktonic growth in medium amended with 0.2% arabinose, the inducer of the expression for the Pbad promoter on the pMQ72 plasmid (Fig. S9B-C).

Additionally, we assayed the *P. aeruginosa* Δ*pqsL* mutant strain in coculture with a few representative *Streptococcus* spp. from Fig. 1B to determine if the Δ*pqsL* mutant strain has a broad defect in *Streptococcus* growth enhancement (Fig. 3D). We found that for *S. intermedius* 4807, there was a slight but non-significant growth decrease during coculture with *P. aeruginosa* Δ*pqsL*, indicating that the mutant strain is unable to enhance *Streptococcus* growth. Similarly, we saw a non-significant decrease in *S. parasanguinis* 5357 growth in coculture with the Δ*pqsL* mutant compared to wild-type *P. aeruginosa* PA14. We did observe a significant decrease in *S. anginosus* 5535 cells recovered from the coculture with the Δ*pqsL* mutant strain compared to monoculture and coculture conditions with *P. aeruginosa* PA14, indicating that the Δ*pqsL* mutation is contributing to the repression of the growth of *S. anginosus* 5535. We found that both wild-type *P. aeruginosa* PA14 and the Δ*pqsL* mutant strain caused a non-significant reduction in *S. intermedius* 7155 cells recovered from coculture, indicating that both of these *P. aeruginosa* strains may be able to outcompete *S. intermedius* 7155. We saw no significant changes to *P. aeruginosa* growth while in coculture with these representative *Streptococcus* spp. (Fig. S10). Taken together, we can extend our finding to at least one other strain of *Streptococcus*.

### The *P. aeruginosa* Δ*pqsL* mutant likely suppresses *S. sanguinis* SK36 growth by siderophore production via iron sequestration

It has been demonstrated previously that a *pqsL* mutant is deficient in HQNO production and overproduces PQS (49). Exogenous PQS has been demonstrated to chelate iron and to increase the expression of the genes coding for siderophore and phenazine biosynthesis enzymes in *P. aeruginosa* (53–55). Thus, we hypothesized that it was the increased production of PQS and/or increased expression of one or more PQS-regulated genes that caused the observed loss in growth promotion of *S. sanguinis* SK36.

To test our hypothesis, we assayed *S. sanguinis* SK36 in coculture with the Δ*pqsL* mutant strains deficient in production of the virulence factors regulated by PQS: siderophores (Δ*pqsLΔpvdAΔpchE*) and phenazines (Δ*pqsLΔphzA-G1/2*) (51, 53–55). We found that the *P. aeruginosa* Δ*pqsL*Δ*pvdA*Δ*pchE* deletion mutant strain restored *S. sanguinis* SK36 growth enhancement to levels similar to wild-type *P. aeruginosa* PA14 (Fig. 4A) without affecting *P. aeruginosa* growth (Fig. S11). In contrast, the *P. aeruginosa* Δ*pqsL*Δ*phzA-G1/2* mutant did not restore *S. sanguinis* SK36 growth (Fig. 4A, Fig. S11).

**Figure 4.**
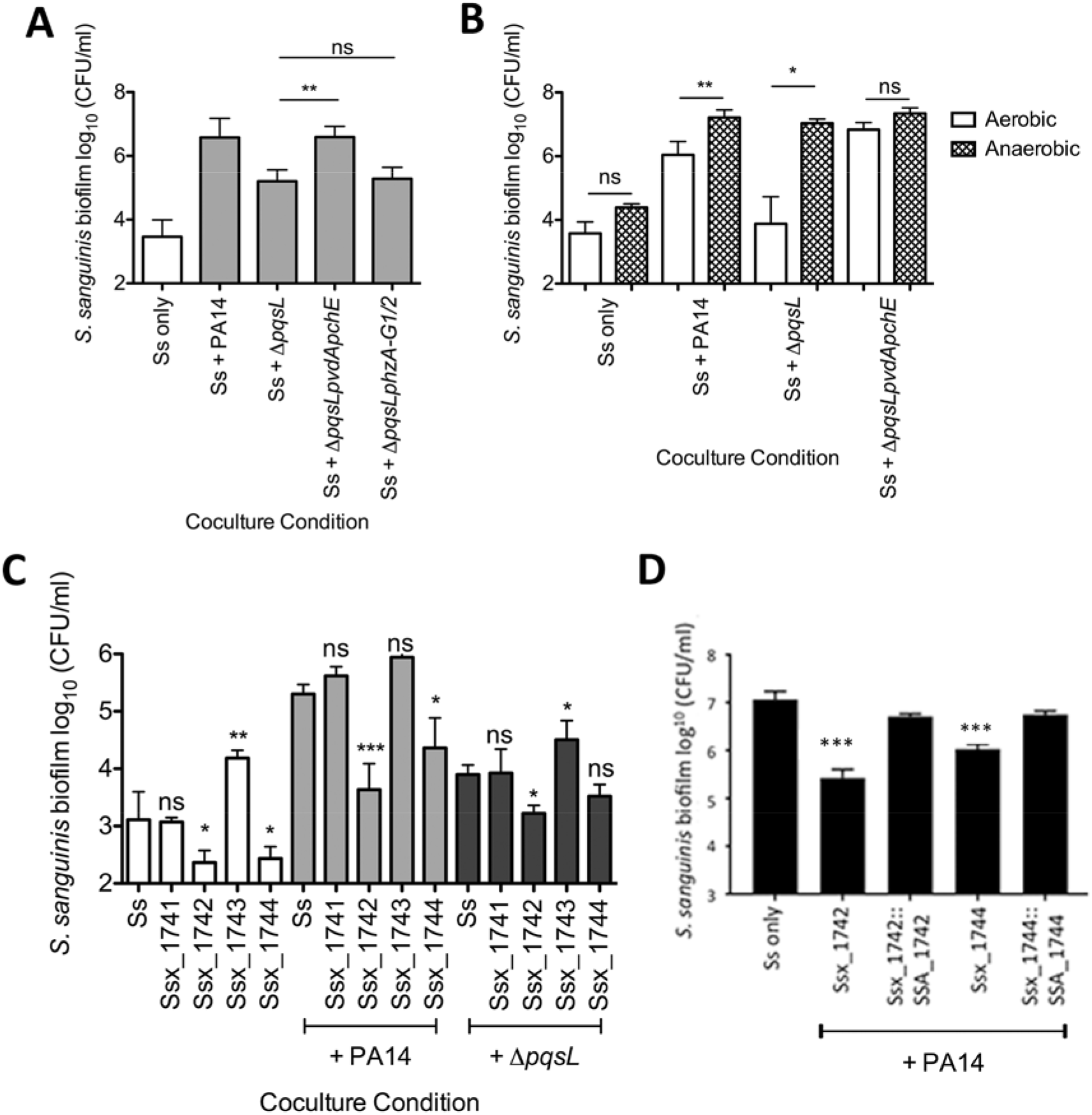
*P. aeruginosa* Δ*pqsL* mutant likely inhibits *Streptococcus* growth by sequestering iron via siderophore production. (A) Coculture of *S. sanguinis* SK36 with *P. aeruginosa* PA14 mutant strains lacking *pqsL*, and siderophore (Δ*pqsLpvdΔpchE*) or phenazine genes (Δ*pqsLphzA-G1/2*). (B) Coculture of *S. sanguinis* SK36 with *P. aeruginosa* PA14 mutant strains in 5% CO_2_ (aerobic) or in anaerobic growth conditions. (C) Coculture of *S. sanguinis* SK36 gene replacement mutants lacking putative iron acquisition genes with *P. aeruginosa* PA14 and the Δ*pqsL* mutant. (D) Complementation assays with the rescued Ssx_1742 and Ssx1744 strains. Each bar represents the average of three biological replicates, each with at least three technical replicates. There was no significant difference between the wild-type *S. sanguinis* SK36 and the two complemented strains. Error bars indicate SD. ns, not significant, ^*^, P < 0.05, ^**^, P < 0.01, and ^***^, P < 0.001 by repeated measures ANOVA with Tukey’s multiple comparisons posttest (A), paired two-tailed student’s t-test (B), repeated measures ANOVA with Dunnett’s multiple comparisons posttest with *S. sanguinis* (Ss) as the control condition (C) and paired two-tailed student’s t-test comparing each complemented and vector control strain, and paired two-tailed student’s t-test with a Bonferroni correction with the WT compared to both complemented strains (D). The corresponding *P. aeruginosa* growth data for these experiments can be found in Supp. Figure S11,13-14.

We performed a complementation experiment to further confirm a role for siderophores in the ability of the Δ*pqsL* mutant to reduce the viability of *Streptococcus.* Specifically, we asked whether complementing the mutation in one of the genes required for siderophore production (*pchE*) would restore the growth defect of the Δ*pqsLΔpvdAΔpchE* mutant to a phenotype similar to that observed for the Δ*pqsL* mutant. We cloned *pchE* into the vector pMQ72, and compared to the vector control strain (ΔpqsLΔpvdAΔpchE/pMQ72), the complemented strain Δ*pqsLΔpvdAΔpchE/* pMQ72-pchE showed a phenotype not significantly different from the Δ*pqsL* mutant (Fig. S12). These data confirmed that it was indeed loss of siderophore production in the Δ*pqsL* mutant background that allowed the Δ*pqsLΔpvdAΔpchE* strain to enhance growth of *S. sanguinis* SK36.

A prediction of this iron sequestration model is that iron supplementation should restore the *Streptococcus* enhancing activity of the *P. aeruginosa* Δ*pqsL* mutant. We added 50μM FeCl_3_ to our minimal medium coculture conditions and saw restoration of *S. sanguinis* SK36 growth enhancement 6 out of 18 times we performed the experiment. We explored this phenotype using a range of FeCl_3_ concentrations from 5μM to 50μM, making a fresh FeCl_3_ solution daily, using buffered media in our assay and when making the FeCl_3_ stock solution, but the phenotype was still variable. We do not fully understand why the iron rescue phenotype was so variable. We measured the iron levels of the medium used in our coculture conditions (MEM) using ICP/MS and showed that the concentration of iron is below the limit of detection (<5 ppb), so it is plausible that the streptococci are iron limited in our coculture conditions.

Next, we tested the idea that coculture in anaerobic conditions would also lead to recovery of the *Streptococcus* growth promotion phenotype in the Δ*pqsL* mutant, because *P. aeruginosa* has been demonstrated to reduce pyoverdine and pyochelin production in anoxic conditions (56). Upon anaerobic coculture in an AnaeroPak-Anaero container with a GasPak satchet, the *P. aeruginosa* Δ*pqsL* mutant significantly enhanced *S. sanguinis* SK36 viability compared to coculture under aerobic conditions (Fig. 4B and Fig. S13A). The level of *S. sanguinis* growth enhancement promoted by the *P. aeruginosa* Δ*pqsL* mutant in anaerobic conditions was equivalent to that observed for the wild-type *P. aeruginosa*. Additionally, if *P. aeruginosa* were enhancing *S. sanguinis* SK36 growth solely through oxygen consumption, we would expect to see enhanced *S. sanguinis* SK36 growth in monoculture under anaerobic conditions, which we do not observe here.

We note that *P. aeruginosa* biofilm and planktonic growth were decreased under the anaerobic growth conditions used in these experiments compared to what we typically observe under aerobic conditions (Fig. S13B-C), which is not surprising given that aerobic respiration is the main means of energy generation for this microbe. Together, these data indicate that wild-type *P. aeruginosa* is contributing to the growth of *S. sanguinis* SK36 via a mechanism independent of oxygen consumption in our coculture system, and are consistent with our hypothesis that reduced siderophore production under anaerobic conditions mitigates the phenotype of the Δ*pqsL* mutant.

### An iron ABC transporter of *S. sanguinis* SK36 participates in competition with *P. aeruginosa*

Our data suggest that one component of the interaction between *P. aeruginosa* and *Streptococcus* spp. is the competition for iron. The genome of *S. sanguinis* SK36 has been sequenced and annotated, and using this information we identified several gene products that, based on their annotation, might be involved in iron uptake. We predicted that if *S. sanguinis* SK36 is indeed competing with *P. aeruginosa* for iron, loss of one or more of these iron uptake system would compromise the ability of *S. sanguinis* SK36 to grow in coculture with *P. aeruginosa*. Given that the *pqsL* mutant likely has an enhanced capacity to scavenge iron as indicated by the restoration of *S. sanguinis* SK36 growth enhancement by the Δ*pqsLΔpvdAΔpchE* mutant, any compromise observed for *S. sanguinis* SK36 iron acquisition mutants should be exacerbated in coculture with the *P. aeruginosa* Δ*pqsL* mutant.

Xu and colleagues reported a mutant library of *S. sanguinis* SK36 wherein nonessential genes are deleted and replaced with a kanamycin resistance cassette (29, 30). Using this library, we examined whether selected *S. sanguinis* SK36 mutant strains lacking genes involved in iron uptake (Table 1) have reduced growth in the presence of wild-type *P. aeruginosa* PA14 or the *P. aeruginosa* Δ*pqsL* strain. We tested *S. sanguinis* SK36 strains carrying mutations in genes coding for iron regulatory proteins, an iron-binding lipoprotein, a ferrichrome-binding protein, and predicted iron-uptake ABC transporters using the coculture assay. We found that all of the mutant strains tested behaved like wild-type *S. sanguinis* SK36 in coculture, except for the *S. sanguinis* SK36 Ssx_1742 and Ssx_1744 mutant strains (Fig. 4C and S14A). The SSA_1742 gene codes for a predicted ferrichrome-binding protein and the SSA_1744 gene codes for a predicted permease protein of an iron compound ABC transporter.

**Table 1:**
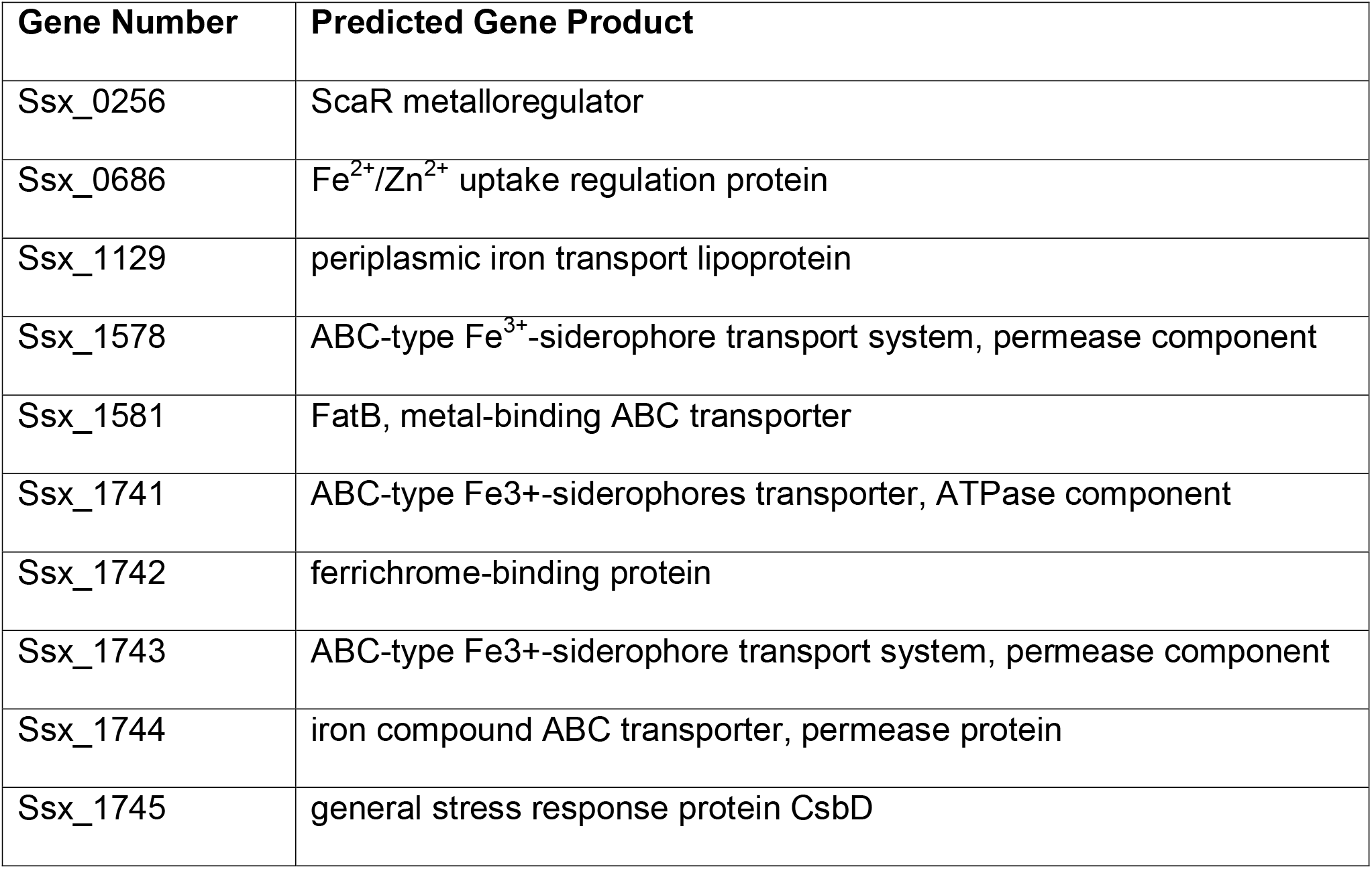
*S. sanguinis* SK36 iron-related gene products.

The *S. sanguinis* SK36 Ssx_1742 and Ssx_1744 mutant strains demonstrated reduced growth in monoculture conditions, 6.5-fold and 6.9-fold respectively, indicating that they may be iron starved in our minimal medium growth conditions (Fig. 4C). This iron starvation phenotype is exacerbated in coculture with wild-type *P. aeruginosa* PA14 with a 44.9-fold reduction in Ssx_1742 cells and a 5.5-fold reduction in Ssx_1744 cells obtained from coculture compared to wild-type *S. sanguinis* SK36 coculture.

To confirm that the observed competition defect was indeed due to the Ssx_1742 and Ssx_1744 mutations, we complemented each of the mutants as described in the Materials and Methods. As shown in Figure 4D, the complemented mutants showed competition phenotypes similar to that observed for wild-type *S. sanguinis* SK36.

We next explored the phenotype of the Ssx_1742 and Ssx_1744 mutants in coculture with the *P. aeruginosa* Δ*pqsL* mutant strain. In coculture with the *P. aeruginosa* Δ*pqsL* mutant strain there is a significant, 4.9-fold reduction in the Ssx_1742 mutant compared to wild-type *S. sanguinis* SK36 in coculture with the Δ*pqsL* mutant. Importantly, coculture of the *S. sanguinis* SK36 Ssx_1744 mutant strain showed no additional, significant growth defect when grown in coculture with the *P. aeruginosa* Δ*pqsL* mutant compared to the wild-type *P. aeruginosa* strain. We take this result to mean that the increased iron sequestration by the Δ*pqsL* mutant is competing for the iron typically transported by the *S. sanguinis* SK36 Ssx_1744-encoded iron ABC transporter; thus loss of Ssx_1744 confers no additional phenotype when cocultured with the *P. aeruginosa* PA14 Δ*pqsL* mutant. Finally, we observed no significant difference in *P. aeruginosa* PA14 and the Δ*pqsL* mutant strain growing in coculture with *S. sanguinis* SK36 mutant strains (Fig. S14B and S14C).

## Discussion

In this study, we sought to characterize a polymicrobial interaction that occurs between *P. aeruginosa* and *Streptococcus* spp. We previously demonstrated that *P. aeruginosa* can suppress *S. constellatus* 7155 biofilms through surfactant production, and that this suppression can be alleviated through treatment with the CF maintenance antibiotic, tobramycin (24). Our current work adds to our understanding of *P. aeruginosa* - *Streptococcus* interactions by demonstrating the widespread ability of multiple *P. aeruginosa* clinical isolates from CF patients and laboratory strains to enhance the growth of multiple species of *Streptococcus*. To better understand the basis of the ability of *P. aeruginosa* to promote the growth of *Streptococcus* spp., we screened *P. aeruginosa* transposon insertion mutants to identify factors that contribute to the ability of *P. aeruginosa* to enhance growth of *S. constellatus* - we identified 46 candidate mutants. Following up on these mutants, we identified only one strain carrying a mutation in the *pqsL* gene that has a consistent, reduced *Streptococcus* spp. growth-enhancement phenotype versus multiple species of *Streptococcus*. Upon further investigation we revealed that this mutant no longer promotes *Streptococcus* growth because the *P. aeruginosa* Δ*pqsL* mutant strain likely actively competes with *Streptococcus* for iron. Loss of PqsL function has been reported to enhance PQS production (49), excess PQS has been demonstrated to enhance siderophore biosynthesis gene transcription (53–55), and PQS-mediated iron sequestration by *P. aeruginosa* has been demonstrated to reduce growth of both Gram-positive and Gram-negative soil bacteria (33). This PQS-mediated growth inhibition of soil bacterial growth can be restored upon addition of 50μM FeCl_3_ (33). Similarly, our data show the ability to restore *Streptococcus* growth by introducing mutations in the siderophore genes to the Δ*pqsL* mutant or by growing the cocultures anaerobically, a growth condition where *P. aeruginosa* is known to reduce pyoverdine and pyochelin production (56). Consistent with these data, we observed that *S. sanguinis* SK36 grew slightly more in the presence of *P. aeruginosa* PA14 Δ*pqsLpvdApchE* mutant than wild-type *P. aeruginosa* PA14. The increased *S. sanguinis* SK36 growth indicates that these microbes are competing for iron during coculture, and that changes to iron uptake can alter the competition between these two microbes. In addition, PQS has been demonstrated to act as both an anti- and pro-oxidant under different conditions (57), and we cannot rule out that the increased PQS production in the Δ*pqsL* mutant impacts production of reactive oxygen species that may be toxic to *Streptococcus* spp.; the fact that growth of the Δ*pqsL* mutant under anaerobic conditions reverses the growth phenotype is consistent with this idea.

An interesting observation from this study was the demonstration of a significant increase in *S. sanguinis* SK36 biofilm growth between an isogenic nonmucoid and mucoid *P. aeruginosa* PAO1 strain. Previous work demonstrated that *S. parasanguinis* is able to use the streptococcal surface adhesin BapA1 to bind alginate produced by mucoid *P. aeruginosa* and enhance *S. parasanguinis* biofilm formation *in vitro*, however *S. gordonii* and *S. sanguinis* SK36 did not demonstrate enhanced biofilm formation (25). Here we demonstrate a significant growth increase of *S. sanguinis* SK36 when in coculture with *P. aeruginosa* PDO300 *mucA22* compared to the isogenic nonmucoid strain. It is possible that *S. sanguinis* SK36 can also bind to alginate. Alternatively, we hypothesize that the growth enhancement induced by the mucoid *P. aeruginosa* strain may be due to decreased rhamnolipid production that has been described in mucoid strains (37), and rhamnolipids (24) and the corresponding relief of rhamnolipid-induced *Streptococcus* killing (24). Furthermore, mucoid strains were shown to produce lower levels of products of the PQS pathway and reduced levels of siderophores (37). Thus, *Streptococcus* spp. may more readily coexist, and perhaps grow to larger numbers, in patients with mucoid *P. aeruginosa*, a question that could be answered by performing a clinical study assessing relative levels of *Streptococcus* spp. as a function of mucoid *P. aeruginosa*. Furthermore, these data indicate that the interactions between *P. aeruginosa* and *Streptococcus* may change over the lifetime of patients with CF as the colonizing *P. aeruginosa* converts to mucoidy.

Oral streptococci have been demonstrated to utilize hydrogen peroxide to inhibit the growth and colonization of competing microorganisms (17, 21, 22, 42), and we hypothesized that *P. aeruginosa* catalase might play a role in enhancing *Streptococcus* growth as catalase has been found in the supernatant of *P. aeruginosa* cultures (39, 40, 43). However, we found no significant defect in *S. sanguinis* SK36 growth enhancement by the *P. aeruginosa ΔkatA, ΔkatB*, and Δ*katAΔkatB* mutants compared to wild-type *P. aeruginosa* PA14, indicating that catalase is not the factor produced by *P. aeruginosa* that is enhancing *Streptococcus* growth. It has been demonstrated that *P. aeruginosa* does not secrete catalase and that it is found in the supernatant due *P. aeruginosa* cell lysis (39) - it may be that catalase found in the supernatant is too dilute to have a positive influence on *Streptococcus* growth in coculture, or that the hydrogen peroxide is not growth limiting to the streptococci in our coculture conditions. Thus, we conclude that *P. aeruginosa* catalase is not influencing *Streptococcus* growth in our model system. It is also worth noting that anaerobic coculture was not sufficient to enhance *S. sanguinis* SK36 monoculture growth to the same levels achieved during coculture with *P. aeruginosa* PA14 in aerobic conditions. These data indicate that *P. aeruginosa-mediated* growth enhancement of streptococci cannot be explained by oxygen consumption via *P. aeruginosa*.

To better understand how *S. sanguinis* SK36 might compete with *P. aeruginosa* for iron in iron-limiting conditions, we examined a set of *S. sanguinis* SK36 mutants lacking putative iron uptake systems or regulatory genes. Of the nine mutants tested, only two showed reduced growth of *S. sanguinis* SK36 when in coculture with *P. aeruginosa* PA14: Ssx_1742 lacking a ferrichrome-binding protein and Ssx_1744 lacking the permease protein of an iron-compound ABC transporter. The Ssx_1742 mutant demonstrated a significant growth defect in monoculture, and during coculture with *P. aeruginosa* PA14. The growth defect of the Ssx_1742 mutant was worsened when cocultured with the Δ*pqsL* mutant. Together, these data indicate that the Ssx_1742 mutant strain is unable to compete with *P. aeruginosa* for the limited iron in our coculture conditions, and that the ferrichrome-binding protein encoded by Ssx_1742 is not involved in the competition for this metal with *P. aeruginosa*. In contrast, the Ssx_1744 mutant showed no additional defect when cocultured with the Δ*pqsL* mutant versus the wild-type *P. aeruginosa*. We take this result to mean that the increased production of the siderophores in the Δ*pqsL* mutant is competing for the iron typically transported by the *S. sanguinis* SK36 Ssx_1744-encoded iron ABC transporter; thus loss of Ssx_1744 confers no additional phenotype when cocultured with the *P. aeruginosa* PA14 Δ*pqsL* mutant. These data indicate that the Ssx_1744-encoded iron ABC transporter of *S. sanguinis* SK36 plays a key role in the competition with *P. aeruginosa*.

Our data support a second mechanism whereby *P. aeruginosa* can limit the growth of *Streptococcus* spp. (Figure 5), including SMG, via iron sequestration. We previously reported that *P. aeruginosa* rhamnolipid surfactants could reduce the viability of *S. constellatus. P. aeruginosa* can also influence the biofilm formation of *S. parasanguinis* through alginate production (25) and the growth of *Streptococcus* spp. via a currently undescribed mechanisms (17, 18, 24). Conversely, previous studies investigating interactions between *P. aeruginosa* and *Streptococcus* spp. also showed that *Streptococcus* spp. influences transcription of *P. aeruginosa* virulence genes, including rhamnolipids, elastase, and phenazine biosynthesis genes through AI-2 signaling (19) and an undescribed mechanism (7, 17, 18), and can suppress *P. aeruginosa* growth when they are a primary colonizer through production of H_2_O_2_ (17, 22) and reactive nitrogenous intermediates (21, 22). Thus, this polymicrobial interaction is complex.

**Figure 5.**
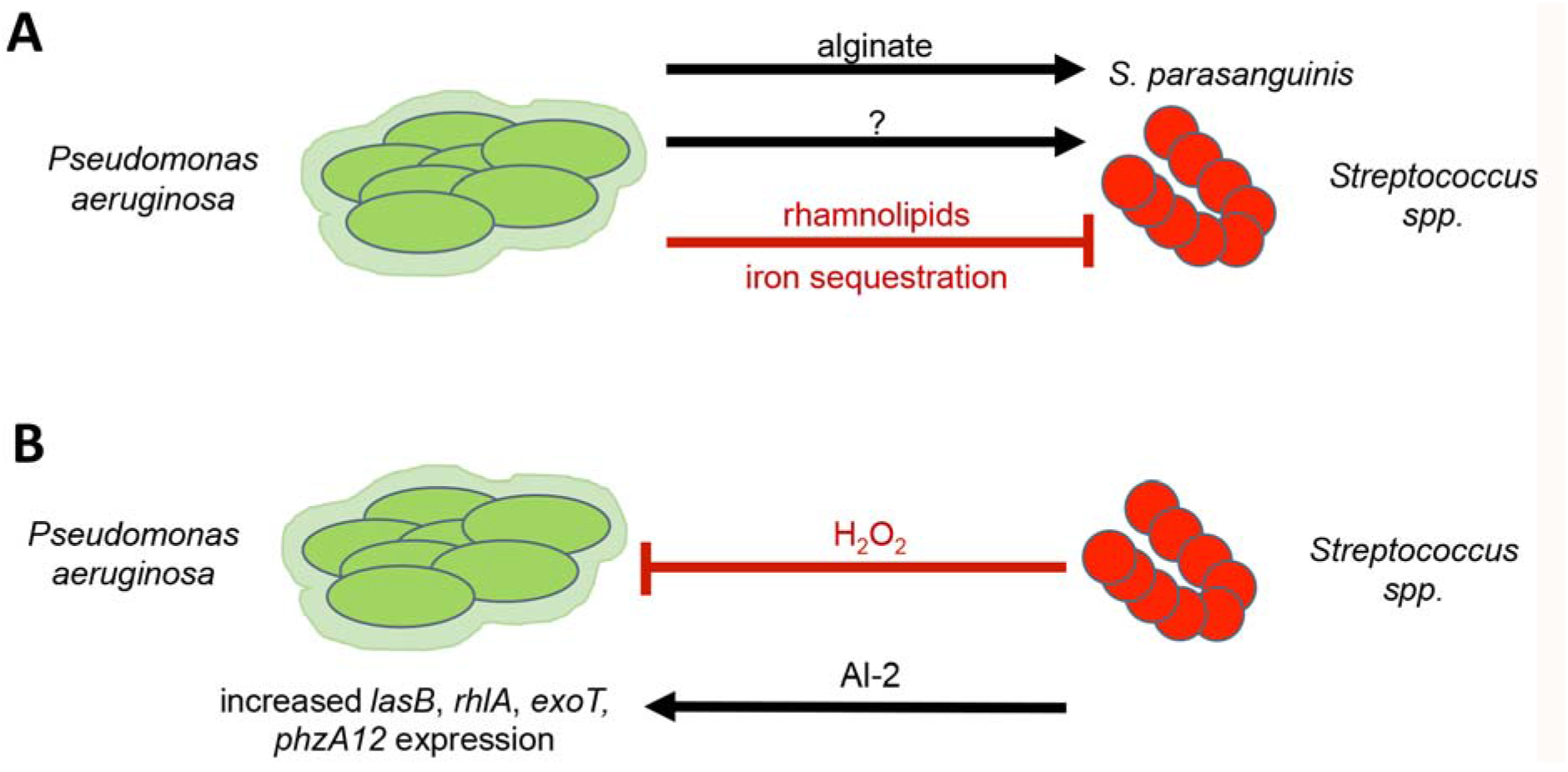
A model for *P. aeruginosa-Streptococcus* interactions. (A-B) *P. aeruginosa* has both positive and negative interactions with *Streptococcus* spp. (A) It has been demonstrated previously that *P. aeruginosa* can enhance *S. parasanguinis* biofilm formation through alginate secretion (25) or inhibit *Streptococcus* growth via rhamonolipid secretion (24). Here we propose a negative interaction wherein iron sequestration by *P. aeruginosa* limits *Streptococcus* spp. growth. *Streptococcus* promoting factors produced by *P. aeruginosa* have not yet been identified (indicated by the question mark). (B) Previous evidence also demonstrates that streptococci can influence *P. aeruginosa* through AI-2 signaling (19), leading to enhanced *lasB, rhlA, exoT*, and *phzA1/2* gene expression, or inhibit *P. aeruginosa* viability through H_2_O_2_ production (17, 21, 22) and subsequent generation of reactive nitrogenous intermediates (21, 22).

Our data also indicate that *P. aeruginosa* can promote the growth of various *Streptococcus* spp., but we do not understand the basis of this growth promotion. We anticipated that the genetic screen described here would likely identify components of such a growth-promoting pathway in *P. aeruginosa;* instead, our screen only identified a single locus apparently involved in a competitive interaction. We suggest two possible explanations for our findings. First, perhaps *P. aeruginosa* determinants that promote *Streptococcus* growth are essential; we think this explanation unlikely, but a formal possibility. More likely is that *P. aeruginosa* has multiple, redundant pathways to boost *Streptococcus* growth. Thus, our genetic approach would be expected to fail to identify such redundant pathways, and alternative strategies to explore *P. aeruginosa-Streptococcus* interactions must be employed in future studies.

Finally, the observations we present here may be of relevance in the CF lung, as many patients are co-colonized by *P. aeruginosa* and *Streptococcus* spp. (9, 58). Analysis of the average available iron in the airway varies markedly between ~0.02 μM in healthy individuals and ~8 μM in patients with CF, and there is a great deal of variability within patients with CF (59, 60). The increased iron in the CF airway is likely due to the reported enhanced levels of extracellular iron in the apical surface liquid of airway cells with a mutation in CFTR (61) and the bleeding into the airway (hemoptysis) associated with this patient population (62). Thus, in CF patients, iron levels in the airway can range from concentrations wherein we might expect direct competition between *P. aeruginosa* and *Streptococcus* for this limited resource, to levels wherein abundant iron would mitigate such competition. Additional studies are necessary to determine if *Streptococcus* spp. are iron limited (or not) in the CF airway, or in sufficiently close proximity to *P. aeruginosa* in the airway (i.e., in mixed microcolonies) to expect direct competition for iron in a local niche.

## Acknowledgements

We thank Dr. Ping Xu for providing his *S. sanguinis* SK36 mutant library and the pVA838 vector, and Dr. Deborah Hogan, Dr. Nicholas Jacobs, and Dr. Dominique Limoli for providing bacterial strains. We thank Dr. Brian Jackson for quantifying the iron concentration in our media with ICP/MS. This work was supported by the Cystic Fibrosis Foundation (OTOOLE16GO), Molecular and Cellular Biology at Dartmouth training grant (T32GM008704), the Munck-Pfefferkorn Fund, and NIH (R37 AI83256-06) to G.A.O and China Scholarship Council (CSC) Grant (201708330005) to K.L.

## Materials and Methods

Bacterial strains and growth conditions. Strains used in this study are listed in Supplemental Table S3. *P. aeruginosa* strains were grown on lysogeny broth (LB) agar or in LB liquid with shaking at 37°C, and where indicated, in the presence of antibiotics at the following concentrations: 25μg/ml gentamicin, 250μg/ml kanamycin, 75μg/ml tetracycline. *Streptococcus* spp. were grown as previously described (24) on tryptic soy agar supplemented with 5% defibrinated sheep’s blood (blood agar) or statically in Todd Hewitt broth supplemented with 0.5% yeast extract (THY) and 20μl/ml oxyrase (Oxyrase, Inc.) at 37°C with 5% CO_2_. *S. sanguinis* SK36 gene replacement mutant strains were grown on blood agar or THY with 500μg/ml kanamycin (30). For antibiotic selection during construction of the *S. sanguinis* SK36 complementation strains, spectinomycin (Spc) was used at a concentration of 100 μg/ml in *E. coli* and 200 μg/ml in *S. sanguinis* SK36.

At the end of each coculture assay, *P. aeruginosa* was grown overnight on *Pseudomonas* Isolation agar (PIA) at 37°C, and *Streptococcus* spp. were grown overnight on blood agar at 37°C anaerobically in AnaeroPak-Anaero containers (Thermo Fisher) or on blood agar supplemented with 10μg/ml neomycin and 10μg/ml polymixin B (*Streptococcus* selection agar) when specified. *Saccharomyces cerevisiae* strain InvSc1 (Invitrogen), was used for homologous recombination to build the pMQ30-*katA*, pMQ-30-katB, and pMQ30-dbpA deletion vector and pMQ72-pqsL complementation vector. InvSc1 was grown as previously described in 1% Bacto yeast extract, 2% Bacto peptone, and 2% dextrose (63). Synthetic defined agar-uracil (4813-065;Qbiogene) was used for InvSc1 selections.

### Species identification of streptococci

*Streptococcus* spp. were isolated at the Dartmouth Hitchcock Medical Center in Lebanon, NH. *Streptococcus* clinical isolates were speciated using 16S rRNA gene sequencing. Genomic DNA (gDNA) was extracted from each strain from overnight cultures using the Gentra Puregene Yeast/Bact. Kit (QIAGEN) followed with 16S-ITS PCR as previously described (64) using the Strep16S-1471F and 6R-IGS primers (listed in Table S4). *Streptococcus oralis, S. mitis*, and *S. pneumoniae* were further differentiated by PCR of a region of the *gdh* gene and sequencing as previously described using the Strep-gdhF and Strep-gdhR primers (listed in Table S4) (64). The Phusion Polymerase PCR protocol (New England Biolabs) was followed for preparing 50μl reactions, and the PCR conditions for the 16S-ITS region were: 98°C for 30s followed by 25 cycles of 98°C for 10s, 61°C for 15s, 72°C for 30s, a final extension at 72°C for 7 minutes. The PCR conditions for amplifying *gdh* were as follows: 98°C for 30s, followed by 30 cycles of 98°C for 10s, 57.9°C for 15s, 72°C for 30s followed by a final extension at 72°C for 7 minutes. The resulting PCR products were imaged on a 1% agarose gel with Sybr Safe (Thermo Fisher Scientific Inc.). The remaining PCR reaction was purified using the QIAquick PCR Purification kit (QIAGEN), and the purified DNA product was sequenced at the Dartmouth Molecular Biology Core Facility using the Applied Biosystems 3730 DNA Analyzer. Sequence results were analyzed using NCBI BLAST for species identification.

### Mixed microbial coculture system

Cocultures were conducted as previously described in the CFBE model system (3, 9, 24, 37) with some modifications. Overnight cultures of *P. aeruginosa* and *Streptococcus* spp. were individually centrifuged at 10,000 x *g* for 3 minutes, the cell pellet was washed with 1.5 ml minimal essential medium (MEM) supplemented with 2 mM L-glutamine (MEM+L-Gln), centrifuged again, and the cell pellet was resuspended in 1.5 ml MEM+L-Gln. The optical density at 600nm (OD_600_) of each culture was determined and the *P. aeruginosa* cultures were adjusted in MEM+L-Gln to an OD600 of 0.05. The *Streptococcus* spp. cultures were adjusted to an OD_600_ of 0.1. *S. sanguinis* SK36 overnight cultures were adjusted to an OD_600_ of 0.1, then further diluted 1:100 in MEM+L-Gln due to the robust growth *S. sanguinis* SK36 exhibits in monoculture. A 1:1 mixture by volume of *P. aeruginosa* and *Streptococcus* spp. was prepared from the adjusted cultures. Three wells of a 96-well plate were inoculated per monoculture and coculture condition with 100uL per well. The culture plates were then incubated statically for 1 hour at 37°C with 5% CO_2_, at which point the unattached planktonic cells were aspirated with a multichannel pipette and replaced with 100μl MEM supplemented with 2mM L-glutamine and 0.4% L-arginine (MEM+L-Gln+L-Arg). The culture plates were incubated statically for an additional 5.5 hours, at which point the supernatant was removed and replaced with 100μl MEM+L-Gln+L-Arg. 0. 4% L-arginine is added to the minimal medium at the 1 and 5.5h medium changes to promote *P. aeruginosa* biofilm formation (65). At 21 hours post-inoculation, planktonic cells were removed to be plated and biofilms were disrupted using a 96 pin replicator in 100μl of MEM+L-Gln. Both planktonic and biofilm cells were 10-fold serially diluted and plated on selective media. PIA plates were grown overnight aerobically, and blood agar plates were grown overnight in AnaeroPak-Anaero containers (Thermo Scientific) with GasPak satchets (BD) to selectively grow *P. aeruginosa* and *Streptococcus* spp., respectively. Following overnight incubation, colonies were counted and the colony forming units (CFU) per milliliter of culture were determined.

### Growth kinetics in mixed microbial coculture system

*P. aeruginosa* PA14 and PAO1 were grown in coculture with *S. sanguinis* SK36 as described above, with one 96-well plate per time point. Six time points were assessed: 0, 3, 5.5, 7.5, 10, and 24 hours. The 0-hour time point corresponds to the initial inoculum. At each time point, the planktonic and biofilm cells from the same wells were serially diluted and plated on PIA and blood agar. Cells were harvested from the 5.5 hour time point plate prior to the 5.5 hour medium exchange.

### Coculture in rich media

*P. aeruginosa* PA14 was grown in coculture with *S. sanguinis* SK36 as described above, but with the following changes: at the 1h media change, the MEM+L-gln was removed and replaced with 100μl LB or THY. 100μl of fresh LB or THY were used at the 5.5h media change as well. This allows all of the culture conditions to originate from the same inocula.

### Construction of *P. aeruginosa* PA14 deletion mutant strains

The pMQ30 vector (Table S3) was used to generate the *P. aeruginosa* PA14 Δ*katA, ΔkatB, ΔkatAΔkatB*, and Δ*dbpA* mutant strains. The pMQ30-katA, pMQ30-katB, and pMQ30-dbpA deletion constructs were built using homologous recombination of the PCR products made with the respective “KO” primers (listed in Table S4) with the Xba1 restriction enzyme-digested pMQ30 in yeast as previously reported (63). Plasmid integrants were isolated on LB agar supplemented with gentimicin and nalidixic acid followed by counterselection on sucrose medium. Deletion mutants were confirmed by PCR with respective “conf.” primers (Table S4), followed by sequencing. Coculture was conducted as described above with the confirmed *P. aeruginosa* PA14 deletion mutant strains.

### Genetic screen

The Ausubel lab created a nonredundant *P. aeruginosa* PA14 transposon library (PA14NR set) in 96-well plate format (65). Initially, a 96 pin replicator was used to transfer inocula from the frozen library to a sterile 96-well plate containing 150μl of LB per well. The plate was then incubated statically for 24 hours at 37°C. *S. constellatus* 7155 frozen aliquots were made from 750μl of overnight culture mixed with 750μl of 40% glycerol. The day of the coculture experiment, frozen *S. constellatus* 7155 aliquots were thawed and 500μl of aliquot were added to 4.5ml THY cultures with 100μl Oxyrase, and were grown for 6-8h at 37°C with 5% CO_2_. The *S. constellatus* 7155 culture was then adjusted to an OD_600_ of 0.05 in MEM+L-Gln, and 100μl of adjusted culture were added to each well of a sterile 96-well plate. The PA14NR set was grown in LB for 24 hr in a 96 well plate format - each well contained a transposon mutant from the *P. aeruginosa* PA14NR set. A 96 pin replicator was then used to transfer 2-3μl of culture from the transposon library plate into the plate containing *S. constellatus* 7155. The coculture plates were then incubated statically for 2 hours at 37°C with 5% CO_2_. After 2 hours, the supernatant and unattached bacteria were aspirated using a multichannel pipette and 100μl MEM+L-Gln with 5μg/ml tobramycin to suppress *P. aeruginosa* PA14 rhamnolipid production were added to each well. The plates were then incubated statically for an additional 20 hours at 37°C with 5% CO_2_. At 22 hours post-inoculation, the 96 pin replicator was used to disrupt the biofilms into the supernatant fraction. The 96 pin replicator was then used to spot culture onto large petri plates containing either PIA or *Streptococcus* selection agar. PIA plates were incubated overnight at 37°C, and *Streptococcus* selection agar plates were incubated for 24 hours at 37°C with 5% CO2. In the initial screen, we identified P. aeruginosa mutants that showed low or undetectable *S. constellatus* 7155 growth. To confirm the phenotype, the candidate *P. aeruginosa* PA14 transposon mutant strain was picked from the PIA plate and grown statically overnight at 37°C in a sterile 96-well plate in 125μl LB. The next morning, 125μl of 40% glycerol was added to each well containing *P. aeruginosa* PA14 candidate mutants, and these “candidate mutant” plates were stored at −80°C for the next round of screening. For the second round of the screen, the coculture process described above was repeated with the plates containing candidate mutants.

If we had clean deletions of the candidate mutants, they were also tested in the assay above. If the clean deletion did not recapitulate the original transposon mutant, that transposon mutant was eliminated from the list of candidate mutants. Table S2 shows the final list of *P. aeruginosa* PA14 transposon insertion mutant strains that yielded low or undetectable *S. constellatus* 7155 growth after rescreening.

We then tested each individual *P. aeruginosa* PA14 transposon mutant in Table S2 in our standard 96-well coculture assay as described above with *S. sanguinis* SK36. The two mutants that yielded consistently low *S. sanguinis* SK36 growth in our standard coculture are in bold in Table S2.

### *P. aeruginosa* Δ*pqsL* complementation

The pMQ72 vector (Table S3) with an arabinose-inducible promoter was used to complement the *P. aeruginosa* PA14 Δ*pqsL* deletion mutant. The pMQ72-pqsL complementation plasmid was built using homologous recombination of the PCR product made with the pqsL comp 3’ and pqsL comp 5’ primers (listed in Table S4) with SacI restriction enzyme-digested pMQ72 in yeast as previously reported (63). *P. aeruginosa* PA14 ΔpqsL/pMQ72-pqsL and *P. aeruginosa* PA14 ΔpqsL/pMQ72-empty vector control strains were cocultured with *S. sanguinis* SK36 as described above with the following changes: at 1 and 5.5h postinoculation, MEM+L-Gln+L-Arg supplemented with L-arabinose at 0% and 0.2% final concentration was added to the medium to induce pMQ72-pqsL gene expression.

### *P. aeruginosa* Δ*pchE* complementation construct

Due to the gene length (4.3 kb) and content of repetitive DNA, the *pchE* gene was amplified in two overlapping PCR fragments using Phusion polymerase (NEB). Fragment 1 was amplified using the primers pchE 5’.2 and pchE int R and fragment 2 was amplified with primers pchE int 1B F (see Table S4 for primer sequences). The resulting PCR fragments were cloned into pMQ72 by homologous recombination in yeast as described above.

### Coculture with ferric chloride

Coculture was conducted as described above, but with the following changes: at 1 and 5.5 hours post-inoculation, supernatants were aspirated with a multichannel pipette, and replaced with MEM+L-Gln+L-Arg, with or without freshly prepared, filter sterilized ferric chloride hexahydrate (ranging from 5 - 50μM).

### Anaerobic coculture

Coculture was conducted as described above, but with the following alterations: once the plates were inoculated, they were incubated in AnaeroPak-Anaero containers with a GasPak satchet. At each medium change (1 hour, 5.5 hours), a new satchet was added to the container to ensure anaerobic coculture conditions. The AnaeroPak container was incubated in the same incubator as the aerobic plate to control for any environmental effects.

### Complementing mutations of *S. sanguinis* SK36

The ORFs of SSA_1742 and SSA_1744 were PCR amplified by PfuUltra II fusion HS DNA polymerase (Agilent Technologies) using primers F-1742-oe/R-1742-oe and F-1744-oe/R-1744-oe, respectively (see Table S4). The PCR productions and an IPTG-inducible plasmid pJFP126 were digested with SphI and/or HindIII, ligated, and electroporated into *E. coli* DH5α (66). Plasmid DNA was purified from DH5α cells using a Qiagen Mini Prep kit (Qiagen). Transformation of *S. sanguinis* strains with each plasmid was carried out as previously described (29). The DNA fragment containing gene expression elements and the *aad9* gene, encoding resistance to Spc, was transferred from the plasmid to the genome of the resulting *S. sanguinis* strains.

## Literature Cited

1. Riordan JR, Rommens JM, Kerem B, Alon N, Rozmahel R, Grzelczak Z, Zielenski J, Lok S, Plavsic N, Chou JL, et al. 1989. Identification of the cystic fibrosis gene: cloning and characterization of complementary DNA. Science 245:1066–73.

2. Elborn JS. 2016. Cystic fibrosis. Lancet 388:2519–2531.

3. Orazi G, O’Toole GA. 2016. *Pseudomonas aeruginosa* alters *Staphylococcus aureus* sensitivity to vancomycin in a biofilm model of Cystic Fibrosis infection. in preparation.

4. Tavernier S, Crabbe A, Hacioglu M, Stuer L, Henry S, Rigole P, Dhondt I, Coenye T. 2017. Community composition determines activity of antibiotics against multispecies biofilms. Antimicrob Agents Chemother 61.

5. Parkins MD, Sibley CD, Surette MG, Rabin HR. 2008. The *Streptococcus milleri* group– –an unrecognized cause of disease in cystic fibrosis: a case series and literature review. Pediatr Pulmonol 43:490–7.

6. Cade A, Denton M, Brownlee KG, Todd N, Conway SP. 1999. Acute bronchopulmonary infection due to *Streptococcus milleri* in a child with cystic fibrosis. Arch Dis Child 80:278–9.

7. Sibley CD, Duan K, Fischer C, Parkins MD, Storey DG, Rabin HR, Surette MG. 2008. Discerning the complexity of community interactions using a *Drosophila* model of polymicrobial infections. PLoS Pathog 4:e1000184.

8. Sibley CD, Sibley KA, Leong TA, Grinwis ME, Parkins MD, Rabin HR, Surette MG. 2010. The *Streptococcus milleri* population of a cystic fibrosis clinic reveals patient specificity and intraspecies diversity. J Clin Microbiol 48:2592–4.

9. Filkins LM, Hampton TH, Gifford AH, Gross MJ, Hogan DA, Sogin ML, Morrison HG, Paster BJ, O’Toole GA. 2012. The prevalence of streptococci and increased polymicrobial diversity associated with cystic fibrosis patient stability. J Bacteriol 194:4709–17.

10. Flight WG, Smith A, Paisey C, Marchesi JR, Bull MJ, Norville PJ, Mutton KJ, Webb AK, Bright-Thomas RJ, Jones AM, Mahenthiralingam E. 2015. Rapid detection of emerging pathogens and loss of microbial diversity associated with severe lung disease in cystic fibrosis. J Clin Microbiol 53:2022–9.

11. Acosta N, Heirali A, Somayaji R, Surette MG, Workentine ML, Sibley CD, Rabin HR, Parkins MD. 2018. Sputum microbiota is predictive of long-term clinical outcomes in young adults with cystic fibrosis. Thorax 73:1016–1025.

12. O’Toole GA. 2018. Cystic fibrosis airway microbiome: Overturning the old, opening the way for the new. J Bacteriol 200.

13. Marshall B, Elbert A, Petren K, Rizvi S, Fink A, Ostrenga J, Sewall A, D. L. 2016. Patient Registry: Annual Data Report 2015. Cyst Fibros Found Patient Regist 194.

14. Rudkjobing VB, Thomsen TR, Alhede M, Kragh KN, Nielsen PH, Johansen UR, Givskov M, Hoiby N, Bjarnsholt T. 2012. The microorganisms in chronically infected end-stage and non-end-stage cystic fibrosis patients. FEMS Immunol Med Microbiol 65:236–44.

15. Sibley CD, Parkins MD, Rabin HR, Duan K, Norgaard JC, Surette MG. 2008. A polymicrobial perspective of pulmonary infections exposes an enigmatic pathogen in cystic fibrosis patients. Proc Natl Acad Sci U S A 105:15070–5.

16. Hogan DA, Willger SD, Dolben EL, Hampton TH, Stanton BA, Morrison HG, Sogin ML, Czum J, Ashare A. 2016. Analysis of lung microbiota in bronchoalveolar lavage, protected brush and sputum samples from subjects with mild-to-moderate Cystic Fibrosis lung disease. PLoS One 11:e0149998.

17. Whiley RA, Fleming EV, Makhija R, Waite RD. 2015. Environment and colonisation sequence are key parameters driving cooperation and competition between *Pseudomonas aeruginosa* cystic fibrosis strains and oral commensal streptococci. PLoS One 10:e0115513.

18. Whiley RA, Sheikh NP, Mushtaq N, Hagi-Pavli E, Personne Y, Javaid D, Waite RD. 2014. Differential potentiation of the virulence of the *Pseudomonas aeruginosa* cystic fibrosis liverpool epidemic strain by oral commensal *Streptococci*. J Infect Dis 209:769–80.

19. Duan K, Dammel C, Stein J, Rabin H, Surette MG. 2003. Modulation of *Pseudomonas aeruginosa* gene expression by host microflora through interspecies communication. Mol Microbiol 50:1477–91.

20. Waite RD, Qureshi MR, Whiley RA. 2017. Modulation of behaviour and virulence of a high alginate expressing *Pseudomonas aeruginosa* strain from cystic fibrosi by oral commensal bacterium *Streptococcus anginosus*. PLoS One 12:e0173741.

21. Scoffield JA, Wu H. 2016. Nitrite reductase is critical for *Pseudomonas aeruginosa* survival during co-infection with the oral commensal *Streptococcus parasanguinis*. Microbiology 162:376–83.

22. Scoffield JA, Wu H. 2015. Oral streptococci and nitrite-mediated interference of *Pseudomonas aeruginosa*. Infect Immun 83:101–7.

23. Waite RD, Qureshi MR, Whiley RA. 2017. Correction: Modulation of behaviour and virulence of a high alginate expressing *Pseudomonas aeruginosa* strain from cystic fibrosis by oral commensal bacterium *Streptococcus anginosus*. PLoS One 12:e0176577.

24. Price KE, Naimie AA, Griffin EF, Bay C, O’Toole GA. 2015. Tobramycin-treated *Pseudomonas aeruginosa* PA14 enhances *Streptococcus constellatus* 7155 biofilm formation in a cystic fibrosis model system. J Bacteriol 198:237–47.

25. Scoffield JA, Duan D, Zhu F, Wu H. 2017. A commensal streptococcus hijacks a *Pseudomonas aeruginosa* exopolysaccharide to promote biofilm formation. PLoS Pathog 13:e1006300.

26. Xu P, Alves JM, Kitten T, Brown A, Chen Z, Ozaki LS, Manque P, Ge X, Serrano MG, Puiu D, Hendricks S, Wang Y, Chaplin MD, Akan D, Paik S, Peterson DL, Macrina FL, Buck GA. 2007. Genome of the opportunistic pathogen *Streptococcus sanguinis*. J Bacteriol 189:3166–75.

27. Stover CK, Pham XQ, Erwin AL, Mizoguchi SD, Warrener P, Hickey MJ, Brinkman FS, Hufnagle WO, Kowalik DJ, Lagrou M, Garber RL, Goltry L, Tolentino E, Westbrock-Wadman S, Yuan Y, Brody LL, Coulter SN, Folger KR, Kas A, Larbig K, Lim R, Smith K, Spencer D, Wong GK, Wu Z, Paulsen IT. 2000. Complete genome sequence of *Pseudomonas aeruginosa* PA01, an opportunistic pathogen. Nature 406:959–64.

28. Winsor GL, Van Rossum T, Lo R, Khaira B, Whiteside MD, Hancock RE, Brinkman FS. 2009. Pseudomonas Genome Database: facilitating user-friendly, comprehensive comparisons of microbial genomes. Nucleic Acids Res 37:D483–8.

29. Chen L, Ge X, Xu P. 2015. Identifying essential *Streptococcus sanguinis* genes using genome-wide deletion mutation. Methods Mol Biol 1279:15–23.

30. Xu P, Ge X, Chen L, Wang X, Dou Y, Xu JZ, Patel JR, Stone V, Trinh M, Evans K, Kitten T, Bonchev D, Buck GA. 2011. Genome-wide essential gene identification in *Streptococcus sanguinis*. Sci Rep 1:125.

31. Jacobs MA, Alwood A, Thaipisuttikul I, Spencer D, Haugen E, Ernst S, Will O, Kaul R, Raymond C, Levy R, Chun-Rong L, Guenthner D, Bovee D, Olson MV, Manoil C. 2003. Comprehensive transposon mutant library of *Pseudomonas aeruginosa*. Proc Natl Acad Sci U S A 100:14339–14344.

32. Maeda Y, Elborn JS, Parkins MD, Reihill J, Goldsmith CE, Coulter WA, Mason C, Millar BC, Dooley JS, Lowery CJ, Ennis M, Rendall JC, Moore JE. 2011. Population structure and characterization of viridans group streptococci (VGS) including *Streptococcus pneumoniae* isolated from adult patients with cystic fibrosis (CF). J Cyst Fibros 10:133–9.

33. Toyofuku M, Nakajima-Kambe T, Uchiyama H, Nomura N. 2010. The effect of a cell-to-cell communication molecule, *Pseudomonas* quinolone signal (PQS), produced by *P. aeruginosa* on other bacterial species. Microbes Environ 25:1–7.

34. Morales DK, Hogan DA. 2010. *Candida albicans* interactions with bacteria in the context of human health and disease. PLoS pathogens 6:e1000886.

35. Chen AI, Dolben EF, Okegbe C, Harty CE, Golub Y, Thao S, Ha DG, Willger SD, O’Toole GA, Harwood CS, Dietrich LE, Hogan DA. 2014. *Candida albicans* ethanol stimulates *Pseudomonas aeruginosa* WspR-controlled biofilm formation as part of a cyclic relationship involving phenazines. PLoS Pathog 10:e1004480.

36. Filkins LM, Graber JA, Olson DG, Dolben EL, Lynd LR, Bhuju S, O’Toole GA. 2015. Coculture of *Staphylococcus aureus* with *Pseudomonas aeruginosa* drives *S. aureus* towards fermentative metabolism and reduced viability in a cystic fibrosis model. J Bacteriol 197:2252–64.

37. Limoli DH, Ivey ML, Filkins LM, Grahl N, Whitfield G, Howell PL, Hogan DA, O’Toole GA, Goldberg JB. 2016. *Pseudomonas aeruginosa* alginate overproduction promotes co-existence with *Staphylococcus aureus* in a model of cystic fibrosis respiratory infections. manuscript in preparation.

38. Brown SM, Howell ML, Vasil ML, Anderson AJ, Hassett DJ. 1995. Cloning and characterization of the *katB* gene of *Pseudomonas aeruginosa* encoding a hydrogen peroxide-inducible catalase: purification of KatB, cellular localization, and demonstration that it is essential for optimal resistance to hydrogen peroxide. J Bacteriol 177:6536–44.

39. Malhotra S, Limoli DH, English AE, Parsek MR, Wozniak DJ. 2018. Mixed communities of mucoid and nonmucoid *Pseudomonas aeruginosa* exhibit enhanced resistance to host antimicrobials. MBio 9: pii: e00275–18.

40. Hassett DJ, Alsabbagh E, Parvatiyar K, Howell ML, Wilmott RW, Ochsner UA. 2000. A protease-resistant catalase, KatA, released upon cell lysis during stationary phase is essential for aerobic survival of a *Pseudomonas aeruginosa oxyR* mutant at low cell densities. J Bacteriol 182:4557–63.

41. Jakubovics NS, Yassin SA, Rickard AH. 2014. Community interactions of oral streptococci. Adv Appl Microbiol 87:43–110.

42. Zhu L, Kreth J. 2012. The role of hydrogen peroxide in environmental adaptation of oral microbial communities. Oxid Med Cell Longev 2012:717843.

43. Lee JS, Heo YJ, Lee JK, Cho YH. 2005. KatA, the major catalase, is critical for osmoprotection and virulence in *Pseudomonas aeruginosa* PA14. Infect Immun 73:4399–403.

44. Liberati NT, Urbach JM, Miyata S, Lee DG, Drenkard E, Wu G, Villanueva J, Wei T, Ausubel FM. 2006. An ordered, nonredundant library of *Pseudomonas aeruginosa* strain PA14 transposon insertion mutants. Proc Natl Acad Sci U S A 103:2833–8.

45. Shajani Z, Sykes MT, Williamson JR. 2011. Assembly of bacterial ribosomes. Annu Rev Biochem 80:501–26.

46. Gentry RC, Childs JJ, Gevorkyan J, Gerasimova YV, Koculi E. 2016. Time course of large ribosomal subunit assembly in *E. coli* cells overexpressing a helicase inactive DbpA protein. RNA 22:1055–64.

47. Fergie N, Bayston R, Pearson JP, Birchall JP. 2004. Is otitis media with effusion a biofilm infection? Clin Otolaryngol Allied Sci 29:38–46.

48. Gallagher LA, McKnight SL, Kuznetsova MS, Pesci EC, Manoil C. 2002. Functions required for extracellular quinolone signaling by *Pseudomonas aeruginosa*. J Bacteriol 184:6472–6480.

49. D’Argenio DA, Calfee MW, Rainey PB, Pesci EC. 2002. Autolysis and autoaggregation in *Pseudomonas aeruginosa* colony morphology mutants. J Bacteriol 184:6481–9.

50. Pesci EC, Milbank JBJ, Pearson JP, McKnight S, Kende AS, Greenberg EP, Iglewski BH. 1999. Quinolone signaling in the cell-to-cell communication system of *Pseudomonas aeruginosa*. Proc Natl Acad Sci USA 96:11229–11234.

51. Maura D, Hazan R, Kitao T, Ballok AE, Rahme LG. 2016. Evidence for direct control of virulence and defense gene circuits by the *Pseudomonas aeruginosa* quorum sensing regulator, MvfR. Sci Rep 6:34083.

52. Wade DS, Calfee MW, Rocha ER, Ling EA, Engstrom E, Coleman JP, Pesci EC. 2005. Regulation of *Pseudomonas* quinolone signal synthesis in *Pseudomonas aeruginosa*. J Bacteriol 187:4372–80.

53. Rampioni G, Falcone M, Heeb S, Frangipani E, Fletcher MP, Dubern JF, Visca P, Leoni L, Camara M, Williams P. 2016. Unravelling the genome-wide contributions of specific 2-alkyl-4-quinolones and PqsE to quorum sensing in *Pseudomonas aeruginosa*. PLoS Pathog 12:e1006029.

54. Diggle SP, Matthijs S, Wright VJ, Fletcher MP, Chhabra SR, Lamont IL, Kong X, Hider RC, Cornelis P, Camara M, Williams P. 2007. The *Pseudomonas aeruginosa* 4-quinolone signal molecules HHQ and PQS play multifunctional roles in quorum sensing and iron entrapment. Chem Biol 14:87–96.

55. Bredenbruch F, Geffers R, Nimtz M, Buer J, Haussler S. 2006. The Pseudomonas aeruginosa quinolone signal (PQS) has an iron-chelating activit Environ Microbiol 8:1318–29.

56. Sonderholm M, Kragh KN, Koren K, Jakobsen TH, Darch SE, Alhede M, Jensen PO, Whiteley M, Kuhl M, Bjarnsholt T. 2017. *Pseudomonas aeruginosa* aggregate formation in an alginate bead model system exhibits in vivo-like characteristics. Appl Environ Microbiol 83.

57. Haussler S, Becker T. 2008. The *Pseudomonas* quinolone signal (PQS) balances life and death in *Pseudomonas aeruginosa* populations. PLoS Pathog 4:e1000166.

58. Filkins LM, O’Toole GA. 2015. Cystic fibrosis lung infections: Polymicrobial, complex, and hard to treat. PLoS Pathog 11:e1005258.

59. Stites SW, Plautz MW, Bailey K, O’Brien-Ladner AR, Wesselius LJ. 1999. Increased concentrations of iron and isoferritins in the lower respiratory tract o patients with stable cystic fibrosis. Am J Respir Crit Care Med 160:796–801.

60. Stites SW, Walters B, O’Brien-Ladner AR, Bailey K, Wesselius LJ. 1998. Increased iron and ferritin content of sputum from patients with cystic fibrosis c chronic bronchitis. Chest 114:814–9.

61. Moreau-Marquis S, Bomberger JM, Anderson GG, Swiatecka-Urban A, Ye S, O’Toole GA, Stanton BA. 2008. The DeltaF508-CFTR mutation results in increased biofilm formation by *Pseudomonas aeruginosa* by increasing iron availability. American journal of physiology: Lung cellular and molecular physiology 295:L25–37.

62. Coss-Bu JA, Sachdeva RC, Bricker JT, Harrison GM, Jefferson LS. 1997. Hemoptysis: a 10-year retrospective study. Pediatrics 100:E7.

63. Shanks RM, Caiazza NC, Hinsa SM, Toutain CM, O’Toole GA. 2006. *Saccharomyces cerevisiae-based* molecular tool kit for manipulation of genes from gram-negative bacteria. Applied and environmental microbiology 72:5027–36.

64. Nielsen XC, Justesen US, Dargis R, Kemp M, Christensen JJ. 2009. Identification of clinically relevant nonhemolytic *Streptococci* on the basis of sequence analysis of 16S-23S intergenic spacer region and partial *gdh* gene. J Clin Microbiol 47:932–9.

65. Anderson GG, Moreau-Marquis S, Stanton BA, O’Toole GA. 2008. In vitro analysis of tobramycin-treated *Pseudomonas aeruginosa* biofilms on cystic fibrosis-derived airway epithelial cells. Infection and immunity 76:1423–33.

66. Rhodes DV, Crump KE, Makhlynets O, Snyder M, Ge X, Xu P, Stubbe J, Kitten T. 2014. Genetic characterization and role in virulence of the ribonucleotide reductases of *Streptococcus sanguinis*. J Biol Chem 289:6273–87.

