## Supplemental Figures for "*Pseudomonas aeruginosa* Can Inhibit Growth of Streptococcal Species via Siderophore Production"

**Figure S1**

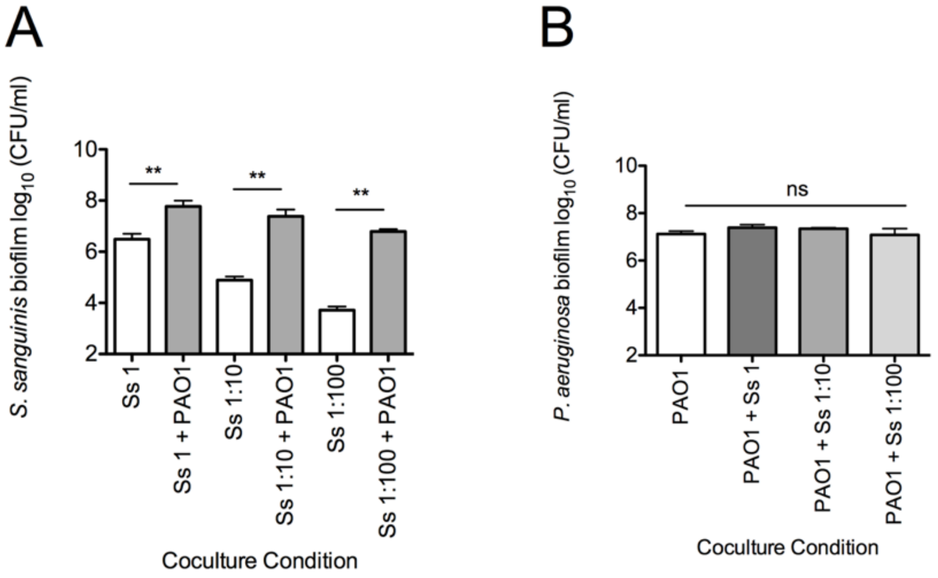


**Figure S1: Growth of *S. sanguinis* SK36 and *P. aeruginosa* PAO1 in coculture.** (A and B) The dilution series of *S. sanguinis* SK36 from an $OD_{600}$ of 0.1 (Ss 1 = undiluted) and the response of each indicated dilution (1:10, 1:100) to coculture with *P. aeruginosa* PAO1 (A) and the corresponding *P. aeruginosa* PAO1 (PAO1) biofilm growth in coculture with each *S. sanguinis* dilution (B). The data shown in Figure 1A and Figure S1 panels A and B are from the same experiments. Each bar represents an average of three biological replicates with three technical replicates. Error bars represent SD. ns, not significant, **, $P < 0.01$ by repeated measures two-tailed student's *t*-test (A) or repeated measures one-way analysis of variance (ANOVA) with Dunnett's post-test using PAO1 as the control (B).

**Figure S2**

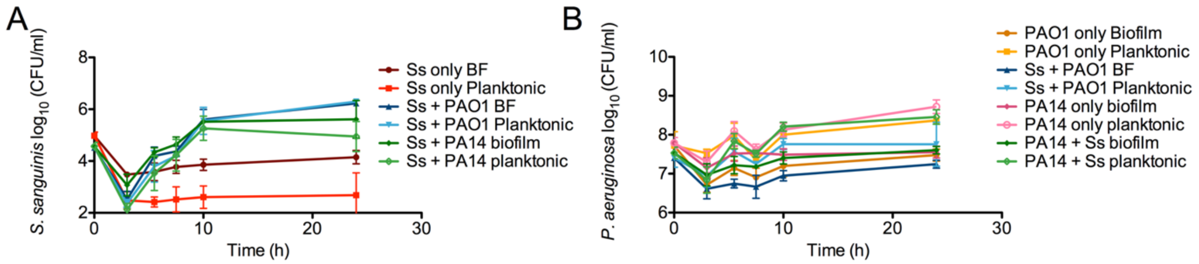


**Figure S2: Coculture growth kinetics of *S. sanguinis* and *P. aeruginosa* PAO1 and PA14 over 24 hours.** (A and B) The growth kinetics of the indicated bacterial species in coculture was investigated over 24 hours, with *S. sanguinis* biofilm and planktonic growth from coculture with *P. aeruginosa* PA14 and PAO1 (A), and the corresponding *P. aeruginosa* PA14 and PAO1 biofilm and planktonic growth (B). The data shown in Figure 1B and Figure S2 panels A and B are from the same experiments. Each time point represents the average of three biological replicates and three technical replicates.

**Figure S3**

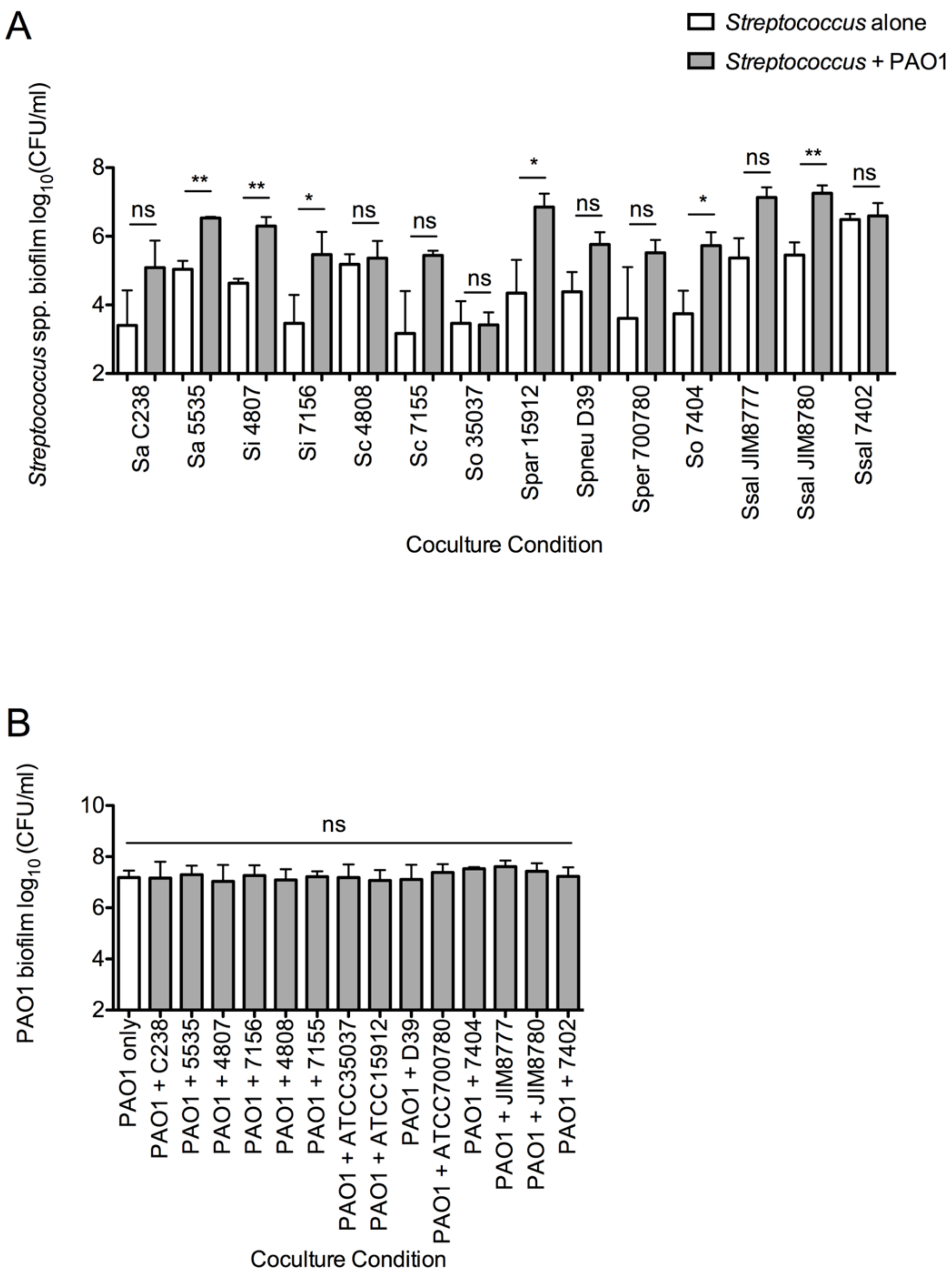
A
Streptococcus alone
Streptococcus + PAO1
Streptococcus spp. biofilm log10(CFU/ml)
8
6
4
2
ns
**
**
*
ns
ns
ns
*
ns
ns
*
ns
**
ns
Sa C238
Sa 5535
Si 4807
Si 7156
Sc 4808
Sc 7155
So 35037
Spar 15912
Spneu D39
Sper 700780
So 7404
Ssal JIM8777
Ssal JIM8780
Ssal 7402
Coculture Condition
B
PAO1 biofilm log10(CFU/ml)
10
8
6
4
2
ns
PAO1 only
PAO1 + C238
PAO1 + 5535
PAO1 + 4807
PAO1 + 7156
PAO1 + 4808
PAO1 + 7155
PAO1 + ATCC35037
PAO1 + ATCC15912
PAO1 + D39
PAO1 + ATCC700780
PAO1 + 7404
PAO1 + JIM8777
PAO1 + JIM8780
PAO1 + 7402
Coculture Condition

**Figure S3: All oral *Streptococcus spp.* tested in coculture with *P. aeruginosa* PAO1 and the corresponding *P. aeruginosa* growth.** (A) Coculture biofilm growth of every oral *Streptococcus spp.* strain tested here with *P. aeruginosa* PAO1. *Streptococcus* spp. are indicated by their strain number and correspond to the following strains: *S. anginosus* C238, *S. anginosus* 5535, *S. intermedius* 4807, *S. intermedius* 7156, *S. constellatus* 4808, *S. constellatus* 7155, *S. oralis* ATCC35037, *S. parasanguinis* ATCC15912, *S. pneumoniae* D39, *S. peroris* ATCC700780, *S. oralis* 7404, *S. salivarius* JIM8777, *S. salivarius* JIM8780, and *S. salivarius* 7402. (B) The growth of *P. aeruginosa* PAO1 in coculture with each oral *Streptococcus spp.* strain tested here. The data shown in Figure S3A and B, and Figure 1C are from the same experiments. Each bar represents the average of three biological replicates with three technical replicates. ns, not significant, *, $P < 0.05$, **, $P < 0.01$ by repeated measures two-tailed student's *t*-test (A) or by repeated measures ANOVA with Dunnett's posttest using PAO1 as the control (B).

### Figure S4

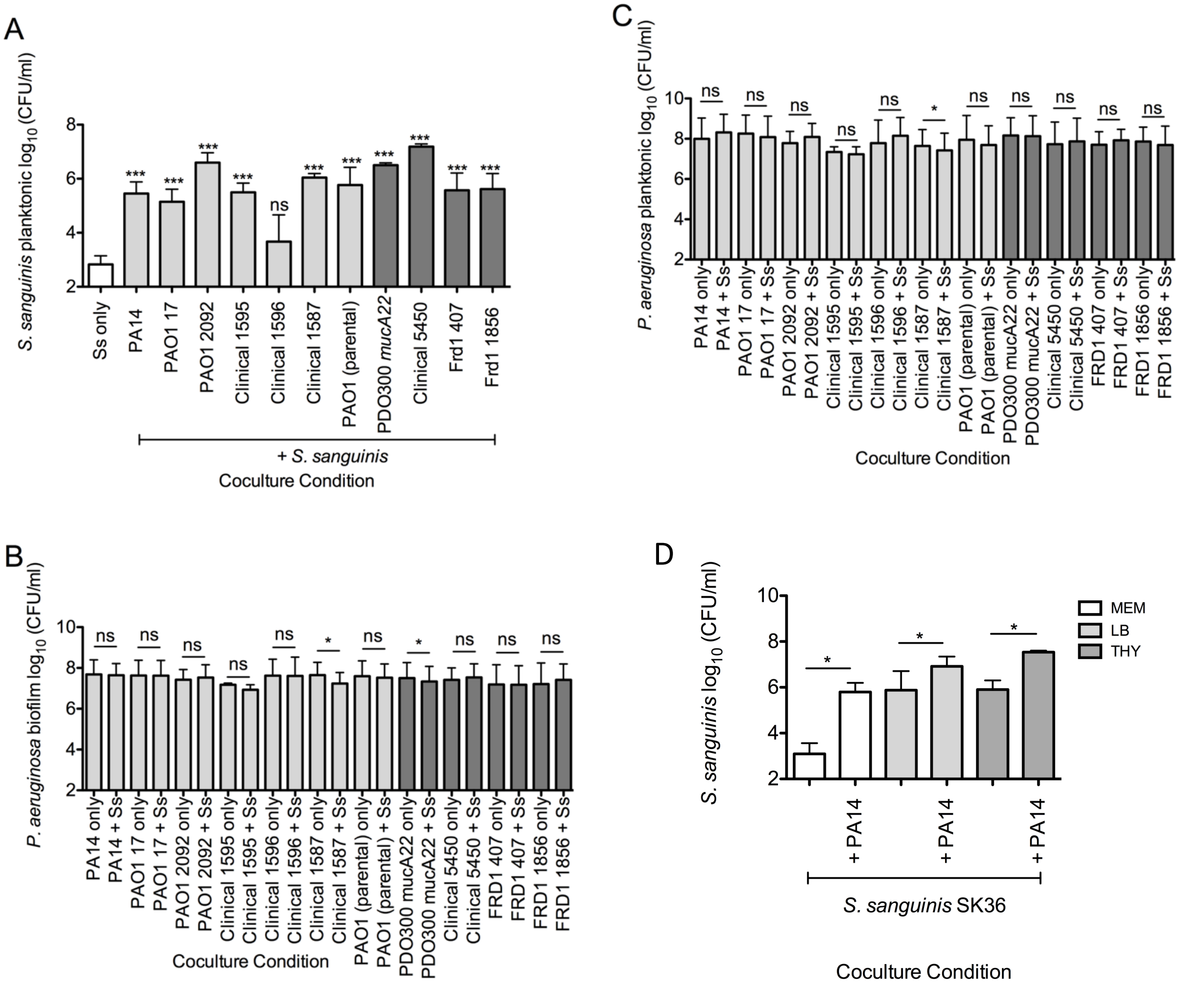


**Figure S4: *S. sanguinis* SK36 planktonic growth, and *P. aeruginosa* biofilm and planktonic growth corresponding to Figure 1.** (A to C) Coculture experiments were conducted with *S. sanguinis* SK36 with different *P. aeruginosa* clinical and laboratory strains to investigate the effects on *S. sanguinis* planktonic growth (A), and *P. aeruginosa* biofilm (B) and *P. aeruginosa* planktonic growth (C). The data shown in Figure 1D and Figure S4 panels A to C are from the same experiments. Bars represent the average of three biological replicates with three technical replicates. Error bars indicate SD. ns, not significant, *, $P < 0.05$, **, $P < 0.01$, ***, $P < 0.001$ by repeated measures ANOVA with Dunnett's posttest for multiple comparisons using Ss only as the control (A), or by repeated measures two-tailed student's *t*-test (B and C). (D) *S. sanguinis* SK36 was tested in coculture with *P. aeruginosa* PA14 in the indicated rich media. The error bars indicate the standard deviation. ns, not significant, *, $P < 0.05$ by paired two-tailed student's *t*-test.

**Figure S5**

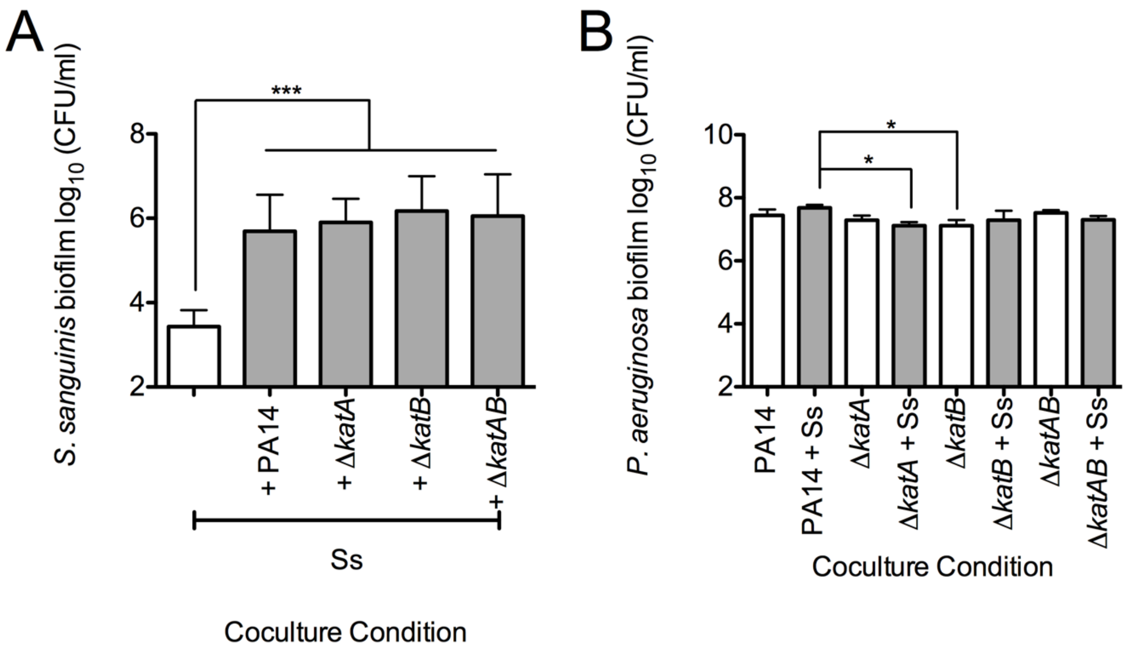


**Figure S5: *P. aeruginosa* catalase mutants do not have a defect in *S. sanguinis* SK36 growth enhancement.** (A) *S. sanguinis* SK36 biofilm growth data from coculture with *P. aeruginosa* PA14 Δ*katA*, Δ*katB*, and Δ*katAB* mutants. (B) Corresponding *P. aeruginosa* biofilm growth data indicates no significant growth defects of *P. aeruginosa* catalase mutants in coculture with *S. sanguinis* SK36. Bars represent the average of three biological replicates with three technical replicates. Error bars indicate SD. ns, not significant by repeated measures ANOVA with Dunnett's post-test for multiple comparisons using Ss only as the positive control (A) and by repeated measures ANOVA with Tukey's post-test for multiple comparisons (B).

**Figure S6**

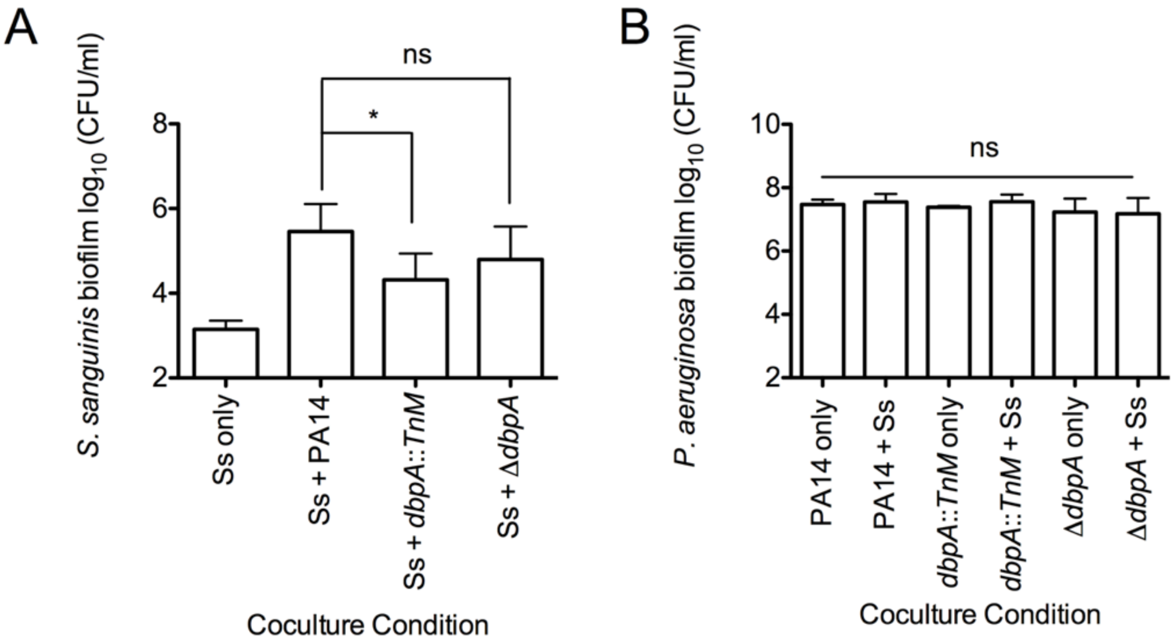


**Figure S6: The *P. aeruginosa* PA14 ∆*dbpA* mutant does not have a defect in *S. sanguinis* SK36 growth enhancement.** (A) *S. sanguinis* SK36 biofilm growth data from coculture with the *P. aeruginosa* PA14 *dbpA*::*TnM* and the ∆*dbpA* mutants. (B) *P. aeruginosa* biofilm growth data from coculture with *S. sanguinis* SK36. Bars represent the average of four biological replicates with three technical replicates. Error bars indicate SD. ns, not significant, *, $P < 0.05$ by repeated measures ANOVA with Dunnett's posttest for multiple comparisons using Ss + PA14 as the control condition.

**Figure S7**

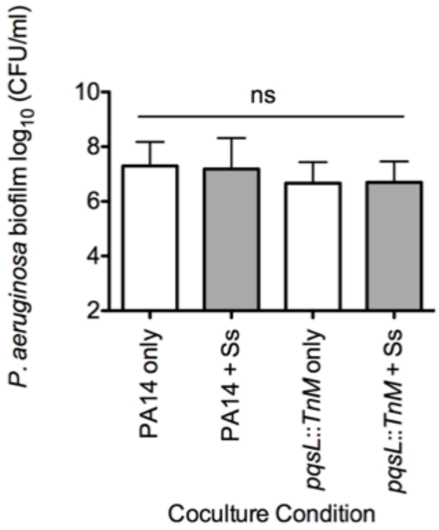


**Figure S7: The *P. aeruginosa* PA14 *pqsL*::*Tn*M mutant does not have a growth defect compared to wild-type *P. aeruginosa* PA14.** *P. aeruginosa* PA14 biofilm growth data from coculture with *S. sanguinis* SK36. The data shown in Figure S7 and Figure 2B are from the same experiments. Bars represent the average of three biological replicates with three technical replicates. Error bars indicate SD. ns, not significant by repeated measures ANOVA with Dunnett's posttest for multiple comparisons using *P. aeruginosa* PA14 only as the control condition.

**Figure S8**

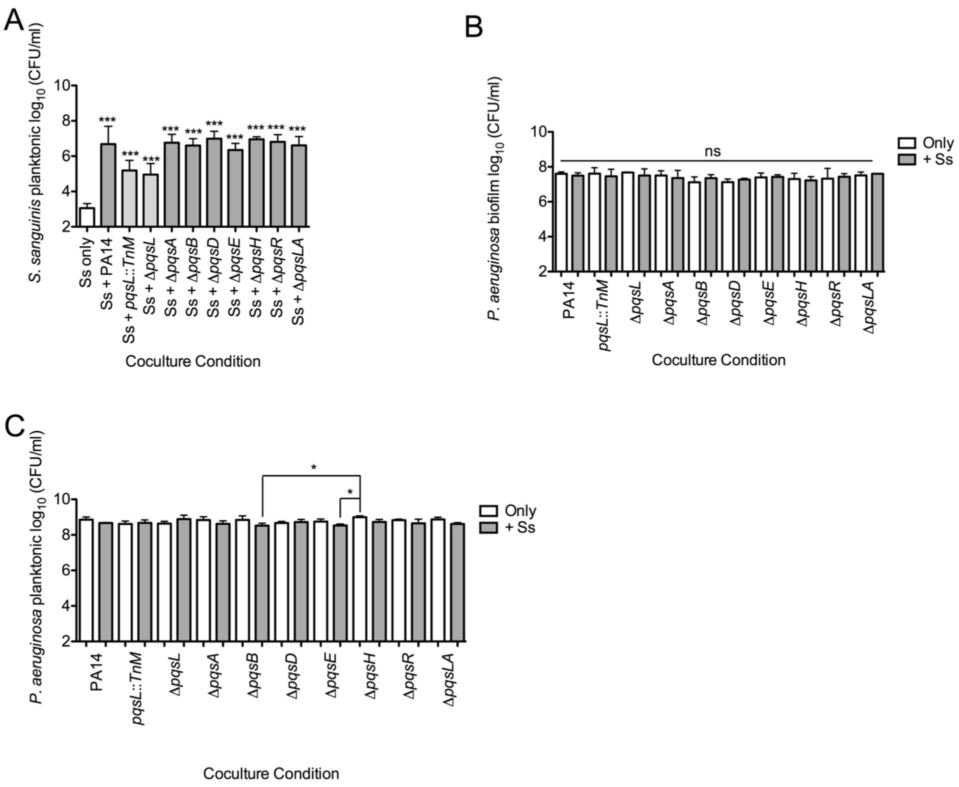


**Figure S8: *S. sanguinis* SK36 planktonic growth in coculture with the *P. aeruginosa* PA14 *pqs* mutant strains, and the corresponding *P. aeruginosa pqs* mutant strain biofilm and planktonic viable counts.** (A to C) *S. sanguinis* SK36 planktonic growth from coculture with *P. aeruginosa* PA14 PQS biosynthetic mutants (A) and the corresponding *P. aeruginosa* PQS biosynthetic mutants' biofilm (B) and planktonic growth data (C). The data shown in Figure 3B and Figure S8 panels A-C are from the same experiments. Bars represent the average of three biological replicates and three technical replicates. Error bars indicate SD. *, $P < 0.05$, ***, $P < 0.001$ by repeated measures ANOVA with Dunnett's posttest for multiple comparisons using Ss only as the control condition (A) or by repeated measures ANOVA with Tukey's posttest for multiple comparisons (B and C).

**Figure S9**

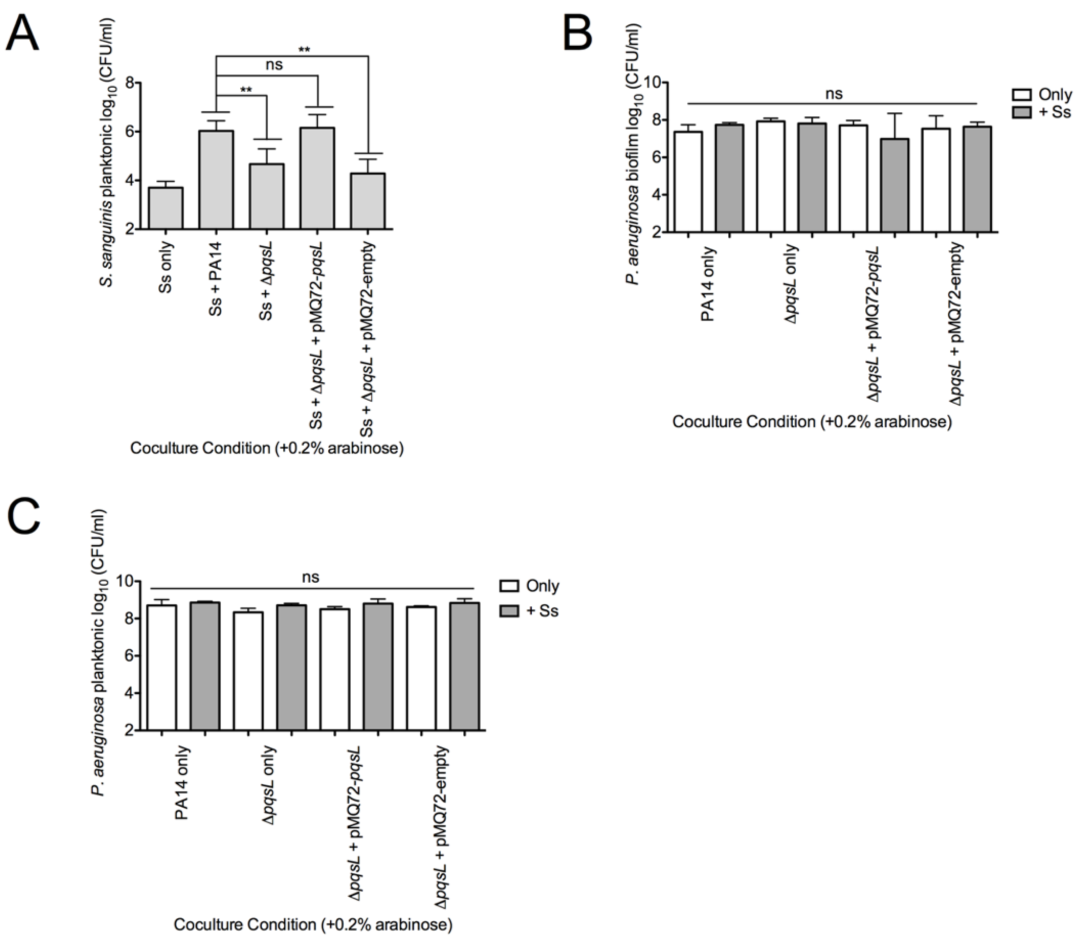


**Figure S9: Quantifying growth from complementation studies.** (A) *S. sanguinis* SK36 planktonic growth data from coculture with wild type *P. aeruginosa* PA14, the Δ*pqsL* mutant, complementation strain and vector control. (B and C) There is no significant difference in *P. aeruginosa* biofilm (B) and planktonic (C) cells recovered from coculture with *S. sanguinis* SK36. The data shown in Figure 3C and Figure S9 panels A-C are from the same experiments. Bars represent three biological replicates with three technical replicates. Error bars indicate SD. ns, not significant, **, $P < 0.01$, by repeated measures ANOVA with Tukey's posttest for multiple comparisons.

**Figure S10**

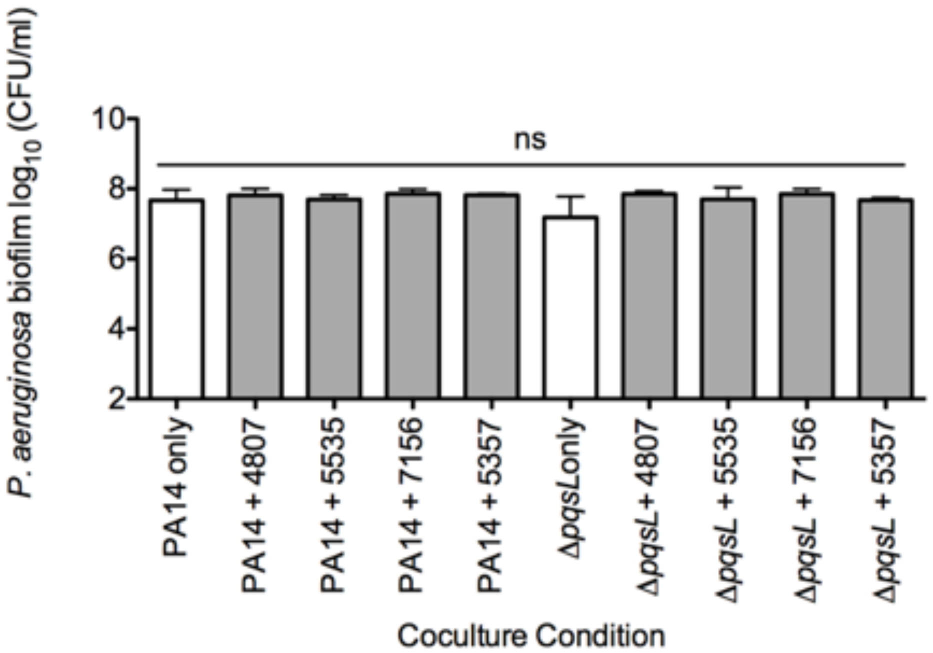


**Figure S10: There is no significant difference between the growth of *P. aeruginosa* strains in the presence of different *Streptococcus* spp.** Growth of the wild-type and mutant *P. aeruginosa* strains in the presence of different *Streptococcus* spp. The data shown in Figures 3D and Figure S10 are from the same experiments. Bars represent three biological replicates with three technical replicates. Error bars indicate SD. ns, not significant by repeated measures ANOVA with Tukey’s posttest for multiple comparisons.

**Figure S11**

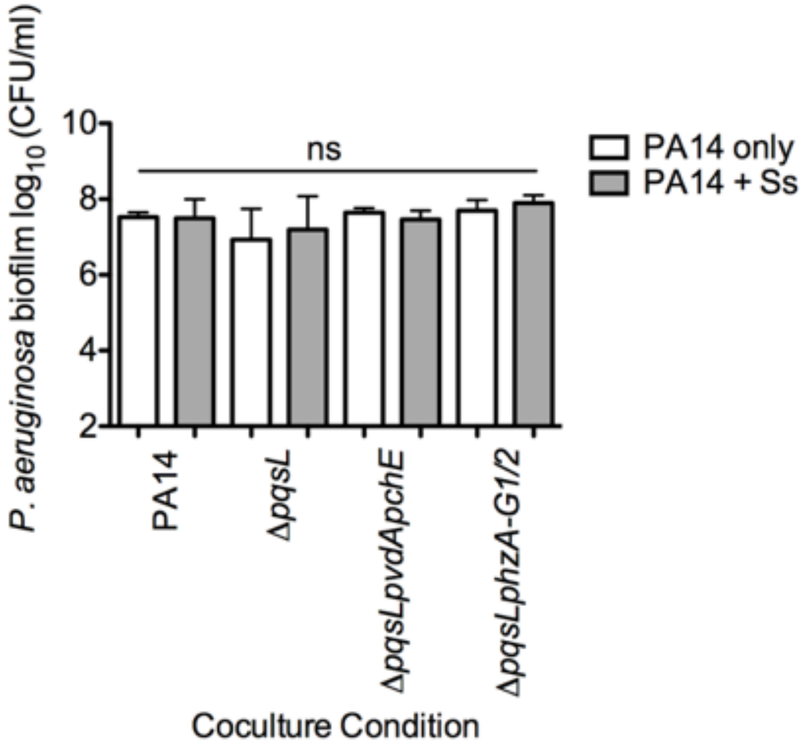


**Figure S11: There is no significant difference between the growth of *P. aeruginosa* PA14 wild type and mutant strains in coculture with *S. sanguinis* SK36.** Growth of the wild type and mutant *P. aeruginosa* PA14 as biofilms from coculture with *S. sanguinis* SK36 (Ss). The data shown in Figures 4A and Figure S11 are from the same experiments. Bars represent three biological replicates with three technical replicates. Error bars indicate SD. ns, not significant by repeated measures ANOVA with Tukey’s posttest for multiple comparisons.

**Figure S12**

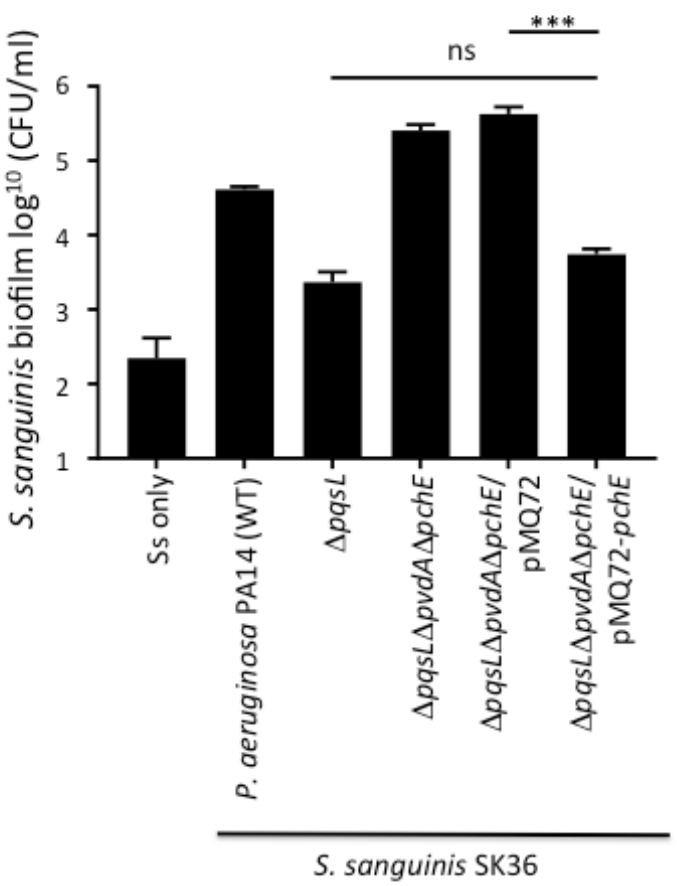


**Figure S12: Complementation analysis.** Shown is the complementation analysis of the Δ*pqsLpvdApchE* mutant in coculture with S. *sanguinis* SK36. Introducing the pMQ72 plasmid carrying the *pchE* gene (pMQ72-*pchE*) but not the vector control (pMQ72) restores the reduced viability of S. *sanguinis* SK36 observed for the Δ*pqsL* mutant. ***, $P < 0.001$ comparing Δ*pqsL*Δ*pvdA*Δ*pchE*/pMQ72 to Δ*pqsL*Δ*pvdA*Δ*pchE*/pMQ72-*pqsE*. Significance was determined with a one-way ANOVA followed by Tukey's multiple comparison. There was no significant difference (ns) between the Δ*pqsL* mutant and the Δ*pqsL*Δ*pvdA*Δ*pchE*/pMQ72-*pqsE* strain using this same test.

**Figure S13**

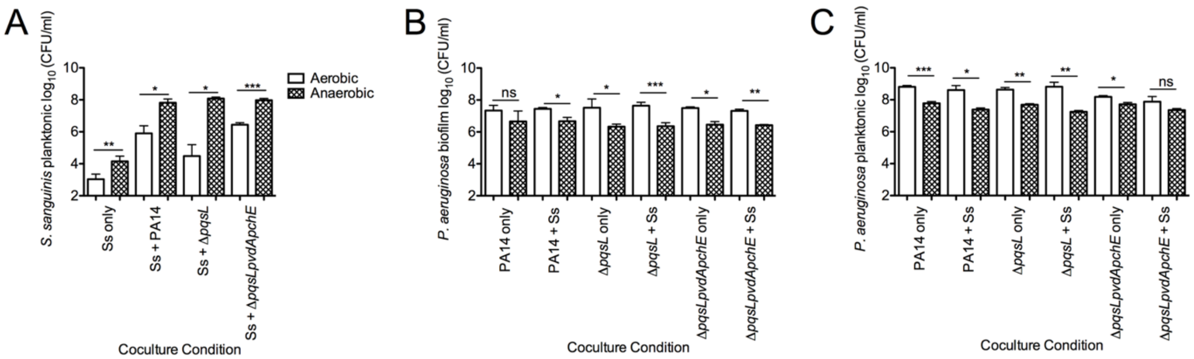


**Figure S13: Growth under anaerobic conditions.** (A) *S. sanguinis* SK36 planktonic growth increases upon coculture with *P. aeruginosa* PA14 in anaerobic conditions. (B and C) *P. aeruginosa* biofilm (B) and planktonic growth in coculture with *S. sanguinis* SK36 in anaerobic conditions are lower (C) compared to *P. aeruginosa* grown in air. The data in Figure 4B and Figure S13 panels A-C are from the same experiments. Bars represent the average of three biological replicates with three technical replicates. Error bars indicate SD. ns, not significant, *, $P < 0.05$, **, $P < 0.01$, ***, $P < 0.001$ by repeated measures two-tailed student's *t*-test.

**Figure S14**

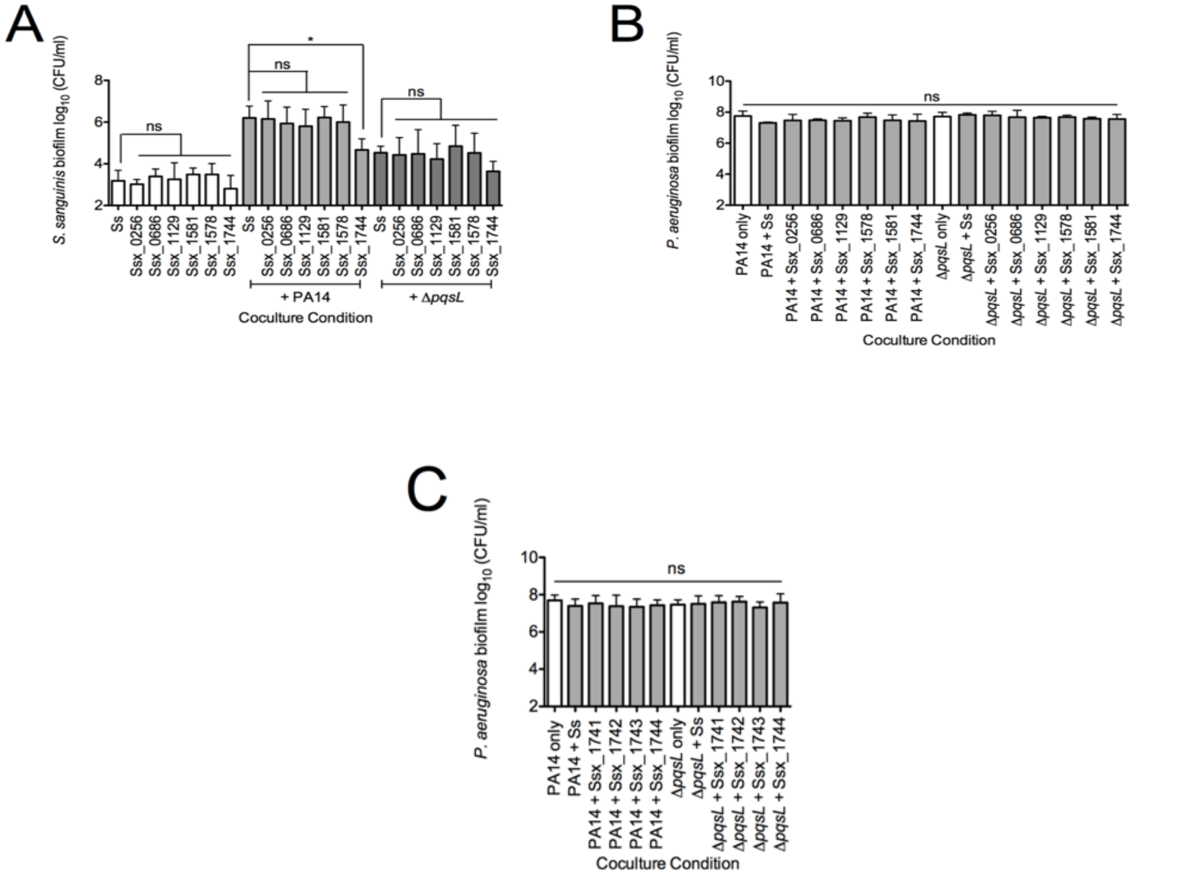


**Figure S14: *S. sanguinis* SK36 iron acquisition mutant strain coculture growth.** (A) *S. sanguinis* SK36 iron acquisition mutants tested in coculture with *P. aeruginosa* PA14 and the Δ*pqsL* mutant strain. (B and C) *P. aeruginosa* biofilm growth from coculture with *S. sanguinis* mutant strains lacking genes for iron acquisition. Figures S14A and B are from the same experiment, and Figures 4C and S14C are from the same experiment. Bars represent the average of three biological replicates with at least three technical replicates. Error bars indicate SD. ns, not significant, *, $P < 0.05$ by one way ANOVA with Dunnett's posttest for multiple comparisons using the Ss condition as the control (A) or with Tukey's posttest for multiple comparisons (B and C).
