## Supplemental Tables for "*Pseudomonas aeruginosa* Can Inhibit Growth of Streptococcal Species via Siderophore Production"

**Supplemental Table S1: Candidate genes tested in coculture with *S. constellatus* 7155 or *S. sanguinis* SK36.**

| **SMC Number** | ***P. aeruginosa* Genotype** | ***S. constellatus* Growth Phenotype** |
|---|---|---|
| SMC1255 | PA14 *algU*::*Tn5phoA*, $Km^r$ | WT[a] |
| SMC1353 | PA14 *rhlA*::$Gm^r$ | WT |
| SMC1503 | PA14 Δ*phnAB* | WT |
| SMC2893 | PA14 Δ*pelA* | WT |
| SMC3718 | PA14 Δ*pilY1* | WT |
| SMC3782 | PA14 Δ*pilA* | WT |
| SMC3809 | PA14 Δ*sadCroeA* | WT |
| SMC5013 | PA14 Δ*pqsA* | WT |
| SMC5017 | PA14 Δ*pqsH* | WT |
| SMC5020 | PA14 Δ$phzA_1$-$G_1$,$A_2$-$G_2$ | WT |
| SMC5021 | PA14 Δ*lasR* | WT |
| SMC5023 | PA14 Δ*lasR rhlR::*$Tet^r$ | WT |
| SMC6215 | PA14 Δ*pvdApchE* | WT |
| 04_3_H3 | PA14 *katA*::*Tn* | WT |
| 05_02_F6 | PA14 *lasB*::*Tn* | WT |
| 05_3_A2 | PA14 *rpoS*::*Tn* | WT |
| DH2528 | PA14 Δ*anr* | WT |
| **SMC Number** | ***P. aeruginosa* Genotype** | ***S. sanguinis* SK36 Growth Phenotype** |
| SMC7814 | PA14 Δ*katA* | WT |
| SMC7815 | PA14 Δ*katB* | WT |
| SMC7842 | PA14 Δ*katA*Δ*katB* | WT |

[a]WT indicates growth enhancement to a level similar to that of *S. constellatus* 7155 or *S. sanguinis* SK36 cocultured with wild-type *P. aeruginosa* PA14.

**Supplemental Table S2: Transposon mutants isolated from the initial PA14NR mutant library screen.**

| Mutant ID | PAO1 Gene Number | PA14 Gene Number | Gene Product Name / Predicted Function | *S. constellatus* growth |
|---|---|---|---|---|
| 23218 | PA3197 | PA14_22860 | hypothetical protein | NG[a] |
| 23227 | PA1786 | PA14_41480 | NasS | low[b] |
| 23324 | PA4441 | PA14_57690 | hypothetical protein | NG |
| 23502 | PA5145 | PA14_67970 | hypothetical protein | low |
| 23525 | PA5475 | PA14_72260 | hypothetical protein | low |
| 23722 | PA0028 | PA14_00320 | hypothetical protein | low |
| 23755 | | PA14_46610 | hypothetical protein | low |
| 23937 | PA1821 | PA14_40980 | probable enoyl-CoA hydratase/isomerase | NG |
| 24001 | PA3957 | PA14_12680 | probable short-chain dehydrogenase | low |
| 24562 | PA3844 | PA14_14290 | hypothetical protein | NG |
| 24838 | PA5352 | PA14_70650 | hypothetical protein | low |
| 25291 | PA2762 | PA14_28380 | hypothetical protein | NG |
| 26867 | PA0790 | PA14_54030 | hypothetical protein | NG |
| 27009 | | PA14_31320 | hypothetical protein | low |
| 27884 | PA0564 | PA14_07340 | probably transcriptional regulator | NG |
| 28880 | PA4521 | PA14_58660 | AmpE | low |
| 31994 | PA2571 | PA14_30840 | probable two-component sensor | NG |
| 32057 | PA4925 | PA14_65040 | hypothetical protein | low |
| 32105 | PA3199 | PA14_22830 | hypothetical protein | NG |
| 33739 | PA5165 | PA14_68230 | DctB | NG |
| 33916 | PA5134 | PA14_67810 | CtpA | low |
| 36263 | PA0573 | PA14_07440 | hypothetical protein | low |

| | | | | |
|---|---|---|---|---|
| 37388 | PA0870 | PA14_53010 | PhhC | NG |
| 37482 | PA4946 | PA14_65350 | MutL | NG |
| 37691 | PA4130 | PA14_10550 | probable sulfite or nitrite reductase | NG |
| 38519 | PA3818 | PA14_14680 | extragenic suppressor protein SuhB | NG |
| 38696 | PA3164 | PA14_23310 | | NG |
| 41259 | PA4588 | PA14_60710 | GdhA | low |
| 41286 | PA4221 | PA14_09340 | FptA | NG |
| 41761 | PA0401 | PA14_05250 | noncatalytic dihydroorotase-like protein | low |
| 41833 | | PA14_59930 | hypothetical protein | NG |
| **45060** | **PA4190** | **PA14_09700** | **PqsL**[d] | **NG** |
| 46072 | | PA14_62950 | hypothetical protein | NG |
| **47831** | **PA4747** | **PA14_62810** | **DbpA** | **NG** |
| 48558 | PA1081 | PA14_50440 | FlgF | NG |
| 48562 | PA0460 | PA14_06010 | hypothetical protein | NG |
| 48720 | PA3391 | PA14_20230 | NosR | NG |
| 55448 | PA1677 | PA14_42820 | hypothetical protein | NG |
| 55651 | PA1851 | PA14_40570 | hypothetical protein | NG |
| 56291 | | PA14_12830* | GID6525 | NG |
| 38595 | PA0401 | PA14_05250 | noncatalytic dihydroorotase-like protein | low |
| 56738 | PA3050 | PA14_24640 | PyrD | NG |
| 26426 | PA1455 | PA14_45630 | FliA | low |
| 37135 | PA1450 | PA14_45710 | hypothetical protein | NG |
| 39797 | PA5267 | PA14_44890 | HcpA | NG |
| 43037 | | PA14_35740 | TnpA | NG |

[a]NG= No growth of *S. constellatus* 7155 in the original screening conditions. See the Material and Methods for details.

[b]low = Less than 20 CFU of *S. constellatus* 7155 in the original screening conditions. See the Material and Methods for details.

**Bold** = the *pqsL*::*TnM* mutant studied here, and the *dbpA*::*TnM* mutant. Both transposon insertion mutants yielded consistently low *S. sanguinis* SK36 growth in our standard coculture assay.

**Supplemental Table S3: Strains and plasmids used in this study.**

| Strain Number | Strain Description | Source |
|---|---|---|
| | ***P. aeruginosa*** | |
| SMC17 | PAO1 | (1) |
| SMC232 | PA14 (wild type) | (2) |
| SMC407 | FRD1 clinical CF isolate (mucoid) | (3) |
| SMC1255 | PA14 *algU*::*Tn5phoA*, $Km^r$ | D. Hogan |
| SMC1353 | PA14 *rhlA*::$Gm^R$ | (4) |
| SMC1503 | PA14 Δ*phnAB* | (5) |
| SMC1587 | CF sputum clinical isolate (mucoid) | (6) |
| SMC1595 | CF sputum clinical isolate (non-mucoid) | (6) |
| SMC1596 | CF sputum clinical isolate (mucoid) | (6) |
| SMC1856 | FRD1 clinical CF isolate (mucoid) | (3) |
| SMC2092 | PAO1 | (7) |
| SMC2893 | PA14 Δ*pelA* | (8) |
| SMC3718 | PA14 Δ*pilY1* | (9) |
| SMC3782 | PA14 Δ*pilA* | (9) |
| SMC3809 | PA14 Δ*sadCroeA* | (10) |
| SMC5013 | PA14 Δ*pqsA* | D. Hogan |
| SMC5014 | PA14 Δ*pqsB* | (11) |
| SMC5015 | PA14 Δ*pqsD* | (11) |
| SMC5016 | PA14 Δ*pqsE* | (11) |
| SMC5017 | PA14 Δ*pqsH* | (12) |
| SMC5018 | PA14 Δ*pqsR* | (12) |
| SMC5020 | PA14 $\Delta phzA_1\text{-}G_1,A_2\text{-}G_2$ | (11) |
| SMC5021 | PA14 Δ*lasR* | (13) |
| SMC5023 | PA14 Δ*lasR rhlR*::*tetR* | (12) |
| SMC5450 | sputum clinical isolate (mucoid) | (6) |
| SMC6215 | PA14 Δ*pvdApchE* | (14) |
| SMC6216 | PA14 Δ*pqsL* | (15) |

| | | |
|---|---|---|
| SMC6217 | PA14 Δ*pqsLpqsA* | (15) |
| SMC6218 | PA14 Δ*pqsApvdApchE* | (15) |
| SMC6219 | PA14 Δ*pqsLpvdApchE* | (15) |
| SMC6220 | PA14 Δ*pqsLphzA-G1/2* | (15) |
| SMC6231 | PA14 Δ*pqsL* pMQ72-*pqsL* complement, $Gm^r$, arabinose inducible | This study |
| SMC6232 | PA14 Δ*pqsL* pMQ72 empty vector control, $Gm^r$ | This study |
| SMC7768 | PAO1 (from Ohman lab, parental for PDO300) | (1)(16) |
| SMC7769 | PAO1 PDO300 *mucA22* | (17) |
| SMC7814 | PA14 Δ*katA* | This study |
| SMC7815 | PA14 Δ*katB* | This study |
| SMC7816 | PA14 Δ*dbpA* | This study |
| SMC7842 | PA14 Δ*katA*Δ*katB* | This study |
| SMC6236 | PA14 Δ*pqsLpvdApchE* pMQ72, $Gm^r$ | This study |
| SMC8244 | PA14 Δ*pqsLpvdApchE* pMQ72-*pchE* complement, $Gm^r$ | This study |
| | ***Streptococcus spp.*** | |
| SMC4806 | *S. intermedius* clinical isolate | This study |
| SMC4807 | *S. intermedius* clinical isolate | This study |
| SMC4808 | *S. constellatus* clinical isolate | This study |
| SMC5355 | *S. oralis* ATCC# 35037 | ATCC |
| SMC5342 | *S. anginosus* C238 | (18) |
| SMC5357 | *S. parasanguinis* ATCC# 15912 | ATCC |
| SMC5443 | *S. pneumoniae* D39 | (19) |
| SMC5535 | *S. anginosus* clinical isolate | This study |
| SMC5767 | *S. salivarius* JIM8777 | (20) |
| SMC5768 | *S. salivarius* JIM8780 | (20) |
| SMC5957 | *S. peroris* ATCC# 700780 | ATCC |
| SMC7155 | *S. constellatus* | This study |
| SMC7156 | *S. intermedius* | This study |
| SMC7402 | *S. salivarius* | This study |

| | | |
|---|---|---|
| SMC7404 | *S. oralis* | This study |
| SMC7474 | *S. sanguinis* SK36 | (21, 22) |
| Ssx_0256 | *S. sanguinis* SK36 Δ*scaR*::$Km^r$; metalloregulator | (23) |
| Ssx_0686 | *S. sanguinis* SK36 ΔSSA_0686::$Km^r$; $Fe^{2+}$/$Zn^{2+}$ uptake regulation protein | (23) |
| Ssx_1129 | *S. sanguinis* SK36 ΔSSA_1129::$Km^r$; periplasmic iron transport lipoprotein | (23) |
| Ssx_1578 | *S. sanguinis* SK36 ΔSSA_1578::$Km^r$; ABC-type $Fe^{3+}$-siderophore transport system, permease component | (23) |
| Ssx_1581 | *S. sanguinis* SK36 ΔSSA_1581::$Km^r$; metal-binding ABC transporter | (23) |
| Ssx_1742 | *S. sanguinis* SK36 SSA_0169::aad9 Phyper-spank lacZo SSA_1742 lacI; $Spc^r$ | (23) |
| SMC8260 | *S. sanguinis* SK36 SSA_1742::aphA-3 SSA_0169::aad9 Phyper-spank lacZo SSA_1742 lacI; $Kan^r$ $Spc^r$ | This study |
| Ssx_1744 | *S. sanguinis* SK36 SSA_0169::aad9 Phyper-spank lacZo SSA_1744 lacI; $Spc^r$ | (23) |
| SMC8261 | *S. sanguinis* SK36 SSA_1744::aphA-3 SSA_0169::aad9 Phyper-spank lacZo SSA_1744 lacI; $Kan^r$ $Spc^r$ | This study |
| | ***Saccharomyces cerevisiae*** | |
| SMC3458 | *S. cerevisiae* InvSc1 | Invitrogen |
| | **Plasmids** | |
| pMQ30 | Vector for gene deletion; $Gm^r$ | (24) |
| pMQ72 | Vector for arabinose inducible gene expression; $Gm^r$ | (24) |
| pVA838 | *E. coli-Streptococcus* shuttle plasmid; $Cm^r$, $Erm^r$ | (25) |

**Supplemental Table S4: Primers used in this study**

| Primer Name | Sequence |
|---|---|
| pqsL comp 5’ | cgctttttatcgcaactctctactgtttctccataatgacggacaaccatatcgatgtac |
| pqsL comp 3’ | cgccaaaacagccaagcttgcatgcctgcaggtcgactcttcagccgcgcggagcctcca |
| pqsL comp conf 1 | gttcaggcgcacctggtcgatc |
| pqsL comp conf 2 | tttcatcctcatgccctgcgagtcg |
| pqsL comp conf 3 | cggcgcttgcaacgcttcg |
| dpqsL conf 5 | cttccggcttgctggtgtggaa |
| Strep16S-1471F | gtgggatag atg attggggtgaagt |
| 6R-IGS | ggg ttc ccc cat tcg gah at |
| Strep-gdhF | atggacaaaccagcnagytt |
| Strep-gdhR | gcttgaggtcccatrctncc |
| dbpA KO-1 | ccgtccttcttgtagcgatgg |
| dbpA KO-2 | gaccttgaacgagcgccccttcggcagcgaggagaaagc |
| dbpA KO-3 | gctttctcctcgctgccgaaggggcgctcgttcaaggtc |
| dbpA KO-4 | ccatgattacgaattcgagctcggtacccggggatccggtagcggctcaccttgg |
| conf. dbpA delF1 | tgtaaaacgacggccagtgccaagcttgcatgcctgcgaaacgactccgaacgg |
| conf. dbpA delF2 | ccgcatcaggtcacgatcg |
| conf. dbpA delR1 | cctgtccggctaattcaatcg |
| conf. dbpA delR2 | tgttcgagcgaggcctgg |
| katA KO-1 | tgtaaaacgacggccagtgccaagcttgcatgcctgaagaagtccctgaccgaaatc |
| katA KO-2 | cgggtcgaccttgaggaacagagcggcagtggtcaggcg |
| katA KO-3 | cgcctgaccactgccgctctgttcctcaaggtcgacccg |
| katA KO-4 | catgattacgaattcgagctcggtacccggggatcctcctacatcaagctgggccagagc |
| conf. katA delF1 | accgtacgttcggccaactg |
| conf. katA delR1 | ggtcagcagcgtgttgaggt |
| conf. katA delF2 | aaagtggtcgtcacctgagcc |

| | |
|---|---|
| conf. katA delR2 | gagaccctggagatccgctaca |
| katB KO-1 | tgtaaaacgacggccagtgccaagcttgcatgcctggacggcttcttcgcgcttgag |
| katB KO-2 | gcccgtaccgtagtcgctctgtacggacagcgacaggag |
| katB KO-3 | ctcctgtcgctgtccgtacagagcgactacggtacgggc |
| katB KO-4 | ccatgattacgaattcgagctcggtacccggggagccctcgacctcgtagatgcg |
| conf. katB delF1 | caggtggcggtggagaactac |
| conf. katB delR1 | cagcatggcctggtcacc |
| conf. katB delF2 | ccttcggcaagttgctcaggc |
| conf. katB delR2 | ggacagagtttccgtttcggc |
| pchE 5'.2 | atacccgtttttttggggaaggagatatacatAGGGAGCCCCCATGGATCTG[a] |
| pchE int R | CGAGTTCCACAGGCTCACCGCATGCCGCTGGATAGCCTCC[a] |
| pchE int 1B F | CTGGACTTCGATCTATCGGTCTTCGACCTGTTC[a] |
| JS pchE 3' | ttaatctgtatcaggctgaaaatcttctctcatccgTCATAGCACGCCCTCCTCCAG[a] |
| F1-1742 | tcatatcatcatttcaggttcttttt |
| R1-1742 | tgtaatcactccttctcactatttactagttcattttttcaaggccatc |
| F3-1742 | ctattatttaacgggaggaaataagattcaagatggccttgaaaaaatg |
| R3-1742 | tgataatagagctaagaggcagacc |
| F1-1744 | catgacagctatttgattttcgact |
| R1-1744 | tgtaatcactccttctcactatttattatttttttttgctgatacataaggtagag |
| F3-1744 | ctattatttaacgggaggaaataagctctaccttatgtatcagcaaaaa |
| R3-1744 | gagtattatcgtgacttgggcttag |
| F2-erm-com | taaatagtgagaaggagtgattacatgaacaa |
| R2-erm-com | ttatttcctcccgttaaataatag |
| pSpank-R | aattcagaacgctcggttgc |
| F-1742-oe | cccaagcttatgaaaaagtttttatcatttgcca |
| R-1742-oe | acatgcatgctgcaaggaatggcctagttc |
| F-1744-oe | acatgcatgcatgaaagataaaaaacggattcttttg |
| R-1744-oe | acatgcatgcagcttggataaataaaaaacaatcagt |

[a]Uppercase letters in the primer sequence indicate complementarity to the pchE gene and lowercase letters indicate complementarity to the pMQ72 plasmid.

**Literature Cited.**


1. Holloway BW, Morgan AF. 1986. Genome organization in *Pseudomonas*. Ann Rev Microbiol 40:79–105.
2. Rahme LG, Stevens EJ, Wolfort SF, Shao J, Tompkins RG, Ausubelt FM. 1995. Common virulence factors for bacterial pathogenicity in plants and animals. Science (80- ) 268:1899–1902.
3. Ohman DE, Chakrabarty AM. 1981. Genetic mapping of chromosomal determinants for the production of the exopolysaccharide alginate in a *Pseudomonas aeruginosa* cystic fibrosis isolate. Infect Immun 33:142–148.
4. Pukatzki S, Kessin RH, Mekalanos JJ. 2002. The human pathogen *Pseudomonas aeruginosa* utilizes conserved virulence pathways to infect the social amoeba *Dictyostelium discoideum*. Proc Natl Acad Sci 99:3159–3164.
5. Mahajan-Miklos S, Tan MW, Rahme LG, Ausubel FM. 1999. Molecular mechanisms of bacterial virulence elucidated using a *Pseudomonas aeruginosa*-Caenorhabditis elegans pathogenesis model. Cell 96:47–56.
6. Yu Q, Griffin EF, Moreau-Marquis S, Schwartzman JD, Stanton BA, O'Toole GA. 2012. In vitro evaluation of tobramycin and aztreonam versus *Pseudomonas aeruginosa* biofilms on cystic fibrosis-derived human airway epithelial cells. J Antimicrob Chemother 67:2673–2681.
7. Jacobs MA, Alwood A, Thaipisuttikul I, Spencer D, Haugen E, Ernst S, Will O, Kaul R, Raymond C, Levy R, Chun-Rong L, Guenthner D, Bovee D, Olson M V, Manoil C. 2003. Comprehensive transposon mutant library of *Pseudomonas aeruginosa*. Proc Natl Acad Sci 100:14339–14344.
8. Friedman L, Kolter R. 2004. Genes involved in matrix formation in *Pseudomonas aeruginosa* PA14 biofilms. Mol Microbiol 51:675–690.
9. Kuchma SL, Ballok AE, Merritt JH, Hammond JH, Lu W, Rabinowitz JD, O'Toole GA. 2010. Cyclic-di-GMP-mediated repression of swarming motility by *Pseudomonas aeruginosa*: The *pilY1* gene and its impact on surface-associated behaviors. J Bacteriol 192:2950–2964.
10. Merritt JH, Ha D, Cowles KN, Lu W, Morales DK, Rabinowitz J, Gitai Z, O'Toole GA. 2010. Specific control of *Pseudomonas aeruginosa* surface-associated

behaviors by two c-di-GMP diquanylate cyclases. MBio 1:e00183-10.

11. Ha D, Merritt JH, Hampton TH, Hodgkinson JT, Janecek M, Spring DR, Welch M, O'Toole GA. 2011. 2-Heptyl-4-Quinolone, a precursor of the *Pseudomonas* Quinolone Signal molecule, modulates swarming motility in *Pseudomonas aeruginosa*. J Bacteriol 193:6770–6780.
12. Cugini C, Morales DK, Hogan DA. 2010. *Candida albicans*-produced farnesol stimulates *Pseudomonas* Quinolone Signal production in LasR-defective *Pseudomonas aeruginosa* strains. Microbiology 156:3096–3107.
13. Hogan DA, Vik Å, Kolter R. 2004. A *Pseudomonas aeruginosa* quorum-sensing molecule influences *Candida albicans* morphology. Mol Microbiol 54:1212–1223.
14. Wang Y, Wilks JC, Danhorn T, Ramos I, Croal L, Newman DK. 2011. Phenazine-1-carboxylic acid promotes bacterial biofilm development via ferrous iron acquisition. J Bacteriol 193:3606–3617.
15. Filkins LM, Graber JA, Olson DG, Dolben EL, Lynd LR, Bhuju S, O'Toole GA. 2015. Coculture of *Staphylococcus aureus* with *Pseudomonas aeruginosa* Drives *S. aureus* towards fermentative metabolism and reduced viability in a cystic fibrosis model. J Bacteriol 197:2252–2264.
16. Martin DW, Schurr MJ, Mudd MH, Govan JRW, Holloway BW, Deretic V. 1993. Mechanism of conversion to mucoidy in *Pseudomonas aeruginosa* infecting cystic fibrosis patients. Proc Natl Acad Sci 90:8377–8381.
17. Mathee K, Ciofu O, Sternberg C, Lindum PW, Campbell JIA, Jensen P, Johnsen AH, Givskov M, Ohman DE, Molin S, Hoiby N, Kharazmi A. 1999. Mucoid conversion of *Pseudomonas aeruginosa* by hydrogen peroxide: a mechanism for virulence activation in the cystic fibrosis lung. Microbiology 145:1349–1357.
18. Olson AB, Kent H, Sibley CD, Grinwis ME, Mabon P, Ouellette C, Tyson S, Graham M, Tyler SD, Domselaar G Van, Surette MG, Corbett CR. 2013. Phylogenetic relationship and virulence inference of *Streptococcus Anginosus* Group: curated annotation and whole-genome comparative analysis support distinct species designation. BMC Genomics 14:1–23.
19. Avery OT, MacLeod CM, McCarty M. 1944. Studies on the chemical nature of the substance inducing transformation of pneumococcal types: Induction of

transformation by a desoxyribonucleic acid fraction isolated from *Pneumococcus* Type III. J Exp Med 79:137–158.

20. Delorme C, Poyart C, Ehrlich SD, Renault P. 2007. extent of horizontal gene transfer in evolution of *Streptococci* of the salivarius group. J Bacteriol 189:1330–1341.
21. Kilian M, Holmgren K. 1981. Ecology and nature of immunoglobulin a1 protease-producing *Streptococci* in the human oral cavity and pharynx. Infect Immun 31:868–873.
22. Hsu SD, Cisar JO, Sandberg AL, Kilian M. 1994. Adhesive properties of viridans streptoccocal species. Microb Ecol Health Dis 7:125–137.
23. Xu P, Ge X, Chen L, Wang X, Dou Y, Xu JZ, Patel JR, Stone V, Trinh M, Evans K, Kitten T, Bonchev D, Buck GA. 2011. Genome-wide essential gene identification in *Streptococcus sanguinis*. Sci Rep 1:1–9.
24. Shanks RMQ, Caiazza NC, Hinsa SM, Toutain CM, O'Toole GA. 2006. *Saccharomyces cerevisiae*-based molecular tool kit for manipulation of genes from gram-negative bacteria. Appl Environ Microbiol 72:5027–5036.
25. Macrina FL, Tobian JA, Jones KR, Evans RP, Clewel DB. 1982. A cloning vector able to replicate in *Escherichia coli* and *Streptococcus sanguis*. Gene 19:345–353.
